# Raman Spectroscopy Enables Real-Time Identification and Monitoring of Plastic Biodegradation Metabolites

**DOI:** 10.64898/2026.06.18.733202

**Authors:** Sydney N. Pedari, Yaxi Hu, David R. McMullin, Paul Heidarian, Allyson Brady, Daniel S. Grégoire

## Abstract

Managing plastic pollution is challenging because current physical and chemical recycling methods are inefficient and environmentally intensive. Biological recycling approaches have been framed as sustainable alternatives but are challenging to optimize due to a lack of process analytical technologies that provide real time data on microbial plastic metabolism. In this study we used *Piscinibacter sakaiensis* 201-F6, a model bacterium with a well-studied polyethylene terephthalate (PET) metabolism, to validate non-destructive Raman spectroscopy methods to monitor plastic biodegradation by tracking metabolite production. Cells were grown on PET and known metabolites stemming from PET metabolism. Raman spectroscopy was used alongside destructive mass spectrometry techniques to monitor PET metabolite production and uptake under different growth conditions. Although cells grew effectively using PET, Raman spectroscopy did not detect the known PET metabolite terephthalic acid during growth assays. Instead, Raman detected isophthalic acid (IPA), a metabolite not previously associated with PET metabolism whose identity was confirmed with LC-HRMS. Raman spectroscopy was also used alongside thermoanalytical techniques to predict the biodegradability of PET at different crystallinities through the release of IPA. This study frames Raman spectroscopy as a promising tool to study metabolic pathways for plastic recycling and optimize their application *in situ*.

## Introduction

Plastics are essential components of packaging and consumer goods, with estimates suggesting 445 million metric tonnes of plastic were produced in 2025 (Houssini et al., 2025; Statista, 2025). Globally, approximately 9% of plastics are properly recycled and similar estimates have been put forward for Canada, which landfills 82% of its plastic waste (OECD, 2022). Ongoing plastic pollution raises significant human and ecosystem health risks due to microplastic exposure (Y. Chen et al., 2022; Sendra et al., 2021) and leaching of harmful additives into ecosystems (*e.g.,* flame retardants and plasticizers) (Covaci et al., 2003). As such, there is a need to develop more effective plastic management strategies that prioritize human and environmental health.

Although ∼75% of plastics produced are recyclable, current physical and chemical processing methods have several limitations that contribute to low recycling rates globally. Mechanical recycling is a low cost approach to reusing plastics, but it is slow and ineffective at repurposing low purity plastics that contain harmful additives (Noor, 2025). Chemical recycling methods (*e.g.,* pyrolysis and solvolysis) that convert plastic polymers into monomers come with high energy demands and the use of hazardous solvents, respectively (Grigore, 2017; Jeswani et al., 2021; Klotz et al., 2024). Biological methods that rely on microbes have been framed as sustainable alternatives to physical and chemical recycling methods but have yet to find widespread application in the solid waste sector (Sun et al., 2024; Zhang et al., 2023). The lack of adoption stems partially from challenges translating lab scale mechanistic studies on model plastic metabolisms into continuous processes that can recycle plastics at the industrial scale.

One of the seminal studies in the field of plastic biodegradation identified a dedicated hydrolase that could depolymerize polyethylene terephthalate (PET) in the model bacterium *Piscinibacter sakaiensis* 201-F6 (formerly known as *Ideonella sakaiensis* 201-F6) (Yoshida et al., 2016). This original study described a mechanism where cells used PET as a carbon source, metabolized PET into mono-(2-hydroxyethyl) terephthalate (MHET) or bis-(2-hydroxyethyl) terephthalate (BHET), and converted these intermediates into terephthalic acid (TPA) and ethylene glycol (EG) to increase biomass (Yoshida et al., 2016). Since this discovery, numerous studies have demonstrated that physical properties of plastics, environmental conditions, and gene regulation networks affect microbial PET metabolism (Palm et al., 2019; Aer et al., 2022; Sevilla et al., 2023; Burgin et al., 2024; Stevensen et al., 2025; Wallace et al., 2020; Walter et al., 2022; Yoshida et al., 2016). Moreover, hundreds of microorganisms have been identified in the literature with the capacity to metabolize plastics other than PET (Gambarini et al., 2022).

Despite the growing repertoire of plastic degrading microbes, the current mechanistic understanding of different plastic metabolisms varies considerably depending on the experimental techniques used in each study. When studies do provide mechanistic details, they rely on gold standard analytical methods for the detection of metabolites such as gas-chromatography mass spectroscopy (GC-MS) and liquid-chromatography high resolution mass spectroscopy (LC-HRMS). These methods have been critical to providing mechanistic details that can be used to optimize plastic biodegradation strategies. However, these methods rely on intensive sample preparation which affects throughput, destroys samples, and makes real-time bioprocess monitoring practically impossible. Overcoming the translational gaps limiting the use of biological plastic recycling strategies requires higher throughput non-destructive methods that can be implemented into process analytical technologies (PAT) *in situ*.

Raman spectroscopy can address these challenges because it is a high throughput non-destructive method that requires no solvents (Uhl et al., 2025). Raman spectroscopy has been used to characterize polymers in solid waste streams (Allen et al., 1999; Asensio-Montesinos et al., 2020; Marica & Pînzaru, 2023; Peñalver et al., 2023) and has been used as a PAT to optimize biological processes related to carbon cycling and fermentation (Borg et al., 2026). The few studies that use Raman to study microbial plastic metabolism focus on structural and chemical changes in parent polymers such as polyethylene (Kang et al., 2019; Tabatabaei et al., 2023; He et al., 2024), polyurethane (Cregut et al., 2014), polylactic acid (Müller et al., 2025) and PET (Santhosh et al., 2025). In contrast, few studies use Raman spectroscopy for real-time identification and monitoring of plastic metabolites, which are critical endpoints to consider when optimizing bioprocess for plastic recycling.

The objective of this study is to validate the use of Raman spectroscopy as a PAT to monitor PET biodegradation in the model bacterium *P. sakaiensis* 201-F6. To achieve this objective, we used Raman spectroscopy to monitor PET metabolite production and uptake over an eight-week period under the same experimental conditions that led to the discovery of the PET hydrolase in *P. sakaiensis*. We compared Raman spectroscopy’s capacity to detect PET biodegradation metabolites to gold standard mass spectrometry techniques including GC-MS and LC-HRMS. We also demonstrated how Raman spectroscopy can characterize PET crystallinity, a key property controlling plastic biodegradability. We frame Raman spectroscopy as an ideal PAT, which can be used to optimize biological plastic recycling strategies and expand our mechanistic understanding of microbial metabolisms used for plastic reclamation.

## Methods and Materials

### PET substrate preparation and characterization

PET powder was purchased from Magerial Science (Wyoming, United States) and passed through a ∼500-mesh size sieve. PET amorphous film (ES30-SH-000115) and bilateral orientation film (ES30-FM-000200) were purchased from Goodfellow (Cambs, England). The film was cut into small pieces (1 cm x 1 cm) and ground using a tube mill (IKA tube-mill 100) in a dry ice bath at 2,500 rpm for 10 seconds. This was repeated until a fine powder was obtained to increase the surface area to volume ratio for biodegradation experiments.

To account for differences in crystallinity of plastics from different suppliers, both the purchased PET powder and film ground to a powder were analyzed using a differential scanning calorimeter (DSC) (Q2000, TA Instruments). Specifically, 9.15 mg of Goodfellow ground (PET-G), 9.89 mg of Magerial Science (PET-M), 14.68 mg of Goodfellow amorphous film, and 7.37 mg of Goodfellow bilateral orientation film were loaded individually onto the instrument at 25 °C, heated to 300 °C at 10 °C min^−1^, held at 300 °C for one minute, then cooled to 25 °C at 10°C min^−1^. Results were analyzed on an TA Instruments Universal Analysis 200 machine. The crystallinities of plastics were calculated using the following equation:

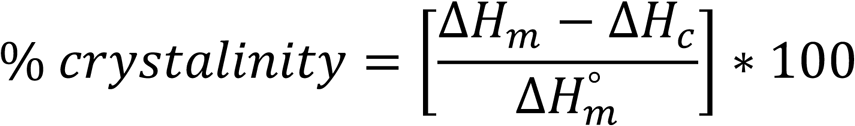

Where Δ*H*_*m*_ is heat of melting, Δ*H*_*c*_ is heat of cold crystallization and Δ*H*^°^*_m_* is the heat of melting of ideal PET crystals at equilibrium (140.1 J g^-1^).

Raman spectroscopy was also used to determine PET crystallinities based on the ratios of key peaks assigned at 1092 cm^-1^ and 1114 cm^-1^ associated with C-C stretching and C-O stretching. These band assignments have been used in previous work, with a higher 1092/1114 cm^-1^ ratio indicating more crystalline material (Stuart, 1996). Briefly, Raman spectra of PET were collected with a Horiba Xplora Plus Raman system (Horiba Scientific Instruments Ltd, Japan). The system was equipped with a 785-nm laser and a Leica microscope with an 100x objective lens (Olympus, NA 0.9, WD 0.21 mm). A laser power of 50% (14 mW) was used to limit the risk of sample burning, and a grating of 600 lines/mm was selected for its broad spectral range to cover both the fingerprint and C-H regions (100 - 2500 cm⁻¹) without sacrificing the high throughput of spectral collection. Each spectrum was acquired with an exposure time of 10 seconds and a replication of 5, taking ∼52 seconds to complete. Raman spectra intensities were normalized using min-max normalization and visualized with RStudio (R version 4.5.2) with the package *plot3D* (Soetaert, 2025).

### Strain cultivation and media preparation

*Piscinibacter sakaiensis* strain 201-F6 was obtained from the DSMZ culture collection (DSM 112585) as a live stab culture. The original stab culture was subsampled and streaked on solid Reasoner’s 2A (R2A) medium (see **Supporting Information** for details on medium composition). Cultures were incubated at 30 °C for 2 days. Following initial growth on a plate, a single colony was transferred into 50 mL of liquid R2A medium in a 250 mL Erlenmeyer flask and incubated at 30 °C while shaking at 175 rpm for 3 days (Thermo Scientific MaxQ 5000 E), at which point cells reached late exponential phase. The culture established in R2A medium was then transferred into a yeast extract-sodium carbonate-vitamins (YSV) medium designed to support PET metabolism by providing essential trace nutrients alongside minimal alternative carbon sources (see **Supporting Information** for details on medium composition).

### PET biodegradation experiments

YSV medium was used for biodegradation assays using PET or known PET biodegradation products (*i.e*., TPA and EG) as starting metabolic substrates. The full factorial design comprising all the abiotic and biotic treatments amended with different carbon substrates is summarized in **Table S1**. Inocula for biodegradation assays were prepared with three-day-old *P. sakaiensis* culture, in 50 mL Falcon tubes that were centrifuged at 2500 ᵡ *g* for 10 minutes. Supernatants were then discarded and the pellet was washed with YSV medium three times.

The different PET powders supplied to cultures were sterilized with 70% ethanol for 24 hours, followed by rinsing with ddH_2_O and drying in a biosafety cabinet (Piedra et al., 2025). The dried PET powder was further sterilized under the UV light in the biosafety cabinet for 20 minutes in line with previous work on PET biodegradation (Piedra et al., 2025). 0.5 g of PET powder acquired from the different suppliers was added to YSV medium for biodegradation experiments. We could not conduct weight-loss analysis due to practical challenges in recovering all the fine PET powder supplied in the flasks. Cultures that were subsampled to monitor growth on PET metabolites received either TPA (1.04 g L^-1^, Sigma-Aldrich, 98%) or EG (1.55 g L^-^1, Sigma-Aldrich, 99%) as carbon substrates. Glucose was used as an alternate carbon source control (1.501 g L^-1^) and culture samples with no defined carbon source were treated as negative controls. Total carbon provided to cells was normalized to 50 mM in all the treatments with defined carbon sources. TPA did not completely dissolve in the medium due to its low solubility in water but was included in experimental treatments to test if it could be metabolized as a sole carbon source.

All cultures were maintained under the same growth conditions for eight weeks, a time frame sufficient to induce detectable PET degradation as reported elsewhere (Wallace et al., 2020). During the eight-week incubation, a 5.7 mL aliquot was subsampled every two weeks to measure cell density using a spectrophotometer (Agilent BioTek Epoch Microplate Reader) at 600 nm and viable plate counts expressed as colony-forming units (cfu/mL). For characterizing the PET metabolites, 5 mL of the 5.7 mL subsamples were centrifuged at 2,500 ᵡ *g* for 10 minutes to remove the cells, and the supernatants were analyzed by LC-HRMS, GC-MS and Raman spectroscopy.

### Analysis of PET-derived metabolites

The 5 mL of aqueous samples obtained from degradation experiments were freeze-dried overnight to remove any remaining water, reconstituted in 2 mL of HR-GC grade methanol (≥99.9%), sonicated for 15 minutes and vortexed for 10 seconds to create homogenous solutions containing metabolite extracts. These extracts were filtered using a 0.22 µm PTFE filter (Thermo Scientific) and transferred into 2 mL amber vials for LC-HRMS, GC-MS, and Raman spectroscopic analyses.

LC-HRMS analyses were performed on an Agilent 1260 Infinity II HPLC system coupled to an Agilent 6230B Time of Flight mass spectrometer (Agilent Technologies). An Eclipse Plus C18 RRHD (2.1×100 mm, 1.8 μm; Agilent Technologies) column was used to resolve the metabolites, with mobile phase A consisting of ddH2O with 0.1% formic acid and mobile phase B consisting of acetonitrile (ACN) with 0.1 % formic acid. The sample injection volume was 2.0 µL, the flow rate was set to a constant 0.4 mL/min, and the gradient chromatographic method is summarized in **Table S2**. The ESI source was operated in negative mode with parameters including a gas temperature of 325 °C, drying time of 10 L/minute, nebulizer of 20 psi, sheath gas temperature of 400 °C at a rate of 12 L/minute, and capillary voltage of 4 kV. The mass-to-charge ratio (*m/z*) range from 50 to 400 *m/z* was scanned for all standards and samples.

Retention times (RT) and *m/z* values of PET metabolite standards were recorded (**Table S3**), which included BHET (Sigma-Aldrich, 96%), MHET (Aladdin Scientific, 98%), TPA (Sigma-Aldrich, 98%) and isophthalic acid (IPA) (Sigma-Aldrich, 99%), an isomer of TPA. BHET, MHET, TPA and IPA were identified based on mass spectra and RT of authentic standards, and the concentration was determined from the signals at *m/z* 165.0188, 209.0450, 165.0188 and 165.0188, respectively, corresponding to their RT (**Figure S1** and **Figure S2**). The calibration curve and regression equations for the PET metabolites are in **Figure S3** and **Table S4,** respectively. Standard curves were fit as second-degree polynomial regression model to improve fits for linear regressions that had R^2^ < 0.9 and are presented in **Table S4** (Agongo et al., 2026).

Data processing was completed on MZmine (Schmid et al., 2023) with the following processing parameters: Mass detection at a noise level of 100, targeted feature detection setting using an *m/z* tolerance of 0.015 or 10 ppm, RT tolerance of 0.3 min and intensity tolerance of 10%, and join aligner settings that included an *m/z* tolerance of 0.02 or 10 ppm and RT tolerance of 0.1 min with equal weights.

Due to the volatility and low molecular size of EG, GC-MS analysis was performed on an Agilent 7890 GC coupled to a 5977B MSD with electron ionization (70 eV) and total ions scanned from *m/z* 29 to 400. A DB-WAX (30 m length x 0.25 mm ID x 0.25 µm film, Agilent Technologies) column was used with helium as the carrier gas (1.0 mL/min). Samples were injected at 250°C in splitless mode. Oven temperature was set to 100 °C for 1 minute, increased to 250 °C at 10°C/min, and held 250 °C for 4 minutes. EG was identified based on mass spectra and RT of authentic standards (Sigma-Aldrich, 99%) and concentrations were determined from the EIC signal at *m/z* 31.0. The RT and calibration curve for EG are in **Table S3, Figure S3** and **Table S4,** respectively.

Raman spectroscopy was applied to analyze controls (medium, abiotic and biotic) and *P. sakaiensis* + PET (G and M) samples at week 2 and week 6 as a rapid and less destructive metabolite identification method. Spectral collection was performed using the same method as described in the previous section for characterizing plastics. A total of 5 µL of each reconstituted sample in methanol was deposited onto a gold-coated glass slide by repeatedly pipetting and drying 1 µL at a time for 5 cycles. Raman spectra of PET metabolite standards (*i.e.*, including MHET, BHET, IPA, TPA and EG) were also collected and are shown in **Figure S4**.

## Results

### Cell growth on PET powders and PET-derived metabolites as carbon sources

*P. sakaiensis* growth assays were conducted in YSV medium amended with PET powders, the PET-derived metabolites (TPA and EG), glucose, or no additional carbon source (**Methods**). These treatments were used to compare growth on PET vs defined carbon sources and demonstrate how Raman spectroscopy could be used for real time monitoring of plastic metabolism.

*P. sakaiensis* cells grown on PET from different suppliers showed similar growth rates based on viable plate counts, which peaked by day 28 at cell densities between 1.1 x 10^7^ to 4.4 x 10^7^ cfu/mL (**Figure 1**). Maximum viable cell densities for cells grown on PET were 2- to 7-fold lower than those obtained by day 28 for cells grown on glucose (ca. 7.7 x 10^7^ to 8.9 x 10^7^ cfu/mL), which aligned with differences observed using OD_600_ (**Figure S5**). Cells provided with EG, one of the products of BHET/MHET metabolism and precursor to the citric acid cycle (Soong et al., 2022), achieved peak viable cell densities by day 28 that were an order of magnitude lower than cultures grown on PET (ca. 3.2 x 10^6^ to 3.9 x 10^6^ cfu/ml) and slightly lower than no carbon controls (ca. 4.8 x 10^6^ to 5.8 x 10^6^ cfu/ml) (**Figure 1**). These trends conflicted with OD_600_ measurements. Cells grown on EG had a turbidity of 0.3 that was comparable to cultures grown on PET or glucose by day 14 prior to declining over the rest of the incubation period while no carbon controls had turbidity ∼0.1 in line with sterile treatments (**Figure S5**). In contrast to EG, cells provided with TPA did not grow based on declining OD_600_ (**Figure S5**) and no detectable viable colonies.

**Figure 1:**
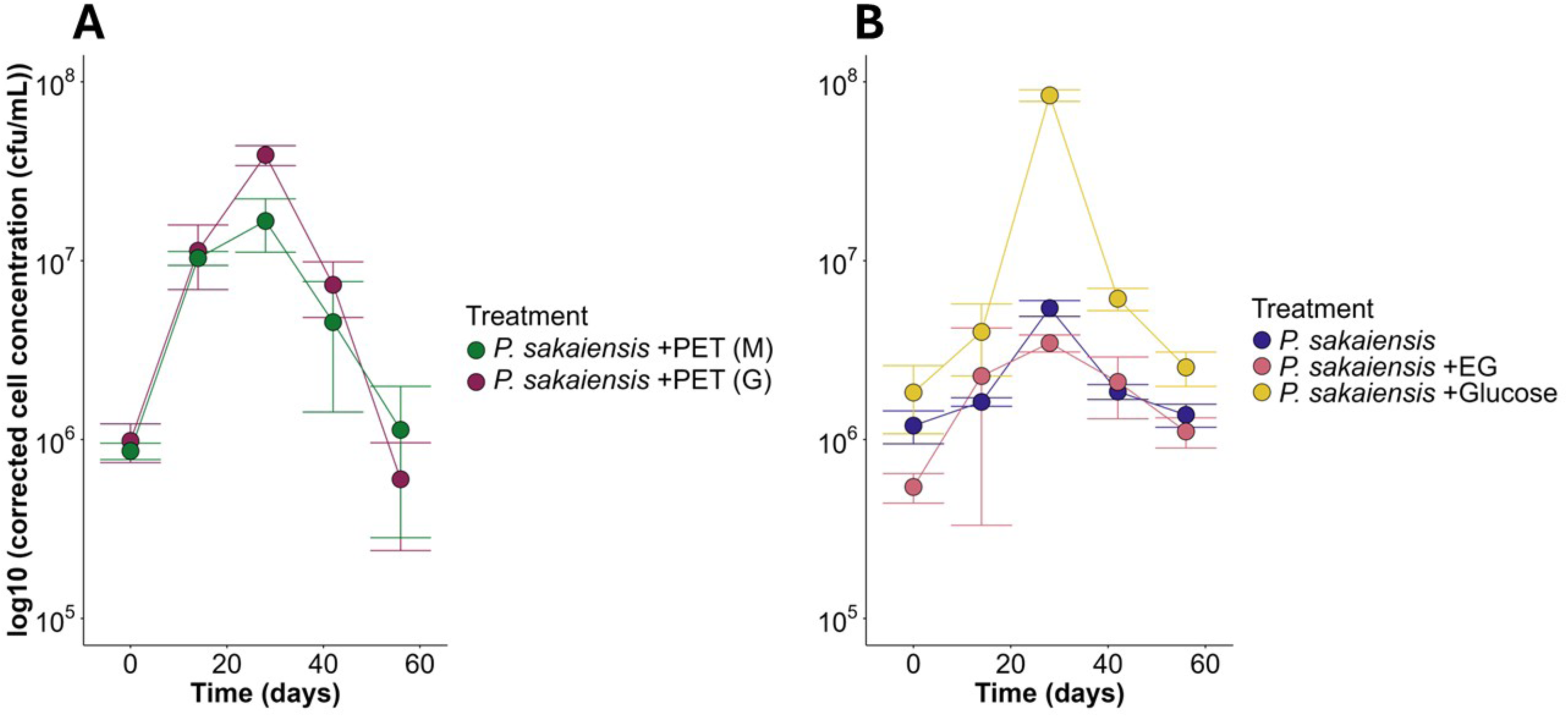
Growth of *Piscinibacter sakaiensis* using the log10 of cell concentrations determined through viable plate counts (cfu/mL ± SD, N=3) throughout the 8-week degradation experiments. (A) Cells grown with two different manufactured polyethylene terephthalates (PET) powders [Goodfellows (PET (G)) and Magerial Science (PET (M))]. (B) Cells grown with ethylene glycol (EG) (50 mM total carbon), glucose (50 mM total carbon), or no carbon at all as controls.

These results show that *P. sakaiensis* grew using PET, glucose, and EG as carbon substrates and retained viability over timeframes representative of biodegradation studies (*i.e.*, > 14 days) (Wallace et al., 2020; Piedra et al., 2025). Despite the discrepancy noted between viable plates counts and OD_600_ measurements for cell grown on EG, these data suggest cells grew moderately on EG. These data also support that cells could not convert EG into biomass as effectively as glucose despite normalizing for total carbon (**Methods**). We attribute the poor growth on TPA to the compound’s low water solubility (17 mg/L at 25°C), which would have reduced its bioavailability to cells.

### Identification and quantification of PET metabolites with MS and Raman spectroscopy

PET-derived metabolites produced by *P. sakaiensis* were analyzed by gold standard methods including LC-HRMS for BHET, MHET and TPA, and GC-MS for EG, to validate whether Raman spectroscopy could detect the same metabolites in a non-destructive manner. LC-HRMS results obtained from negative control samples (*i.e.*, sterile YSV medium and YSV medium with live cells grown in the absence of additional carbon) did not show peaks that aligned with RT for MHET and BHET but did show low intensity peaks for TPA (∼10^4^ vs ∼10^5^ for the lowest concentration in the standard curve of 1 ppm) over the incubation period (**Figure S6**, **Figure S1** and **Table S3**). This suggests that low but non-quantifiable (ca. < 1 ppm) contamination of the TPA monomer occurred due to the experimental setup and sample processing method.

Similar peak areas for TPA were detected in chromatograms for controls where PET was supplied in sterile medium (**Figure S7**, **Figure S1** and **Table S3**). Control samples also showed peak height intensities ranging from ∼10^4^ to 10^5^ for compounds sharing similar *m/z* and RT to BHET (**Figure S7**, **Figure S1** and **Table S3**). For the ion with an *m/z* value of 209.0450 matching to MHET, the EIC showed a peak at an RT earlier than the RT associated with the standard compound (*i.e.*, 5.14 min vs 5.24 min), indicating the presence of a molecule sharing a similar structure to MHET. The low intensities of these BHET and MHET-like peaks compared to their 1 ppm standards suggest leaching of low amounts (ca. < 1 ppm) of each compound occurred in sterile treatments during the incubation. This leaching could be due to incomplete PET polymerization, weathering, or oxidation reactions that released compounds into the medium during the incubation (Shi et al., 2023; Stanica-Ezeanu & Matei, 2021).

Raman spectra collected for abiotic treatments of YSV medium with and without PET shared three strong peaks at 446, 607 and 970 cm^-1^ (**Figure S8**). These bands, which correspond to the N-C-S stretching, glycerol and phosphate monoester groups of phosphorylated proteins and cellular nucleic acid, respectively (Talari et al., 2015), were expected given the composition of the YSV medium used to maintain cells (**Methods**).

PET metabolites (*i.e.,* BHET, MHET and TPA) in samples obtained from cultures grown on PET were analyzed using LC-HRMS and Raman spectroscopy bearing the background signals noted above in mind. No BHET or TPA were detected above the baseline peak intensity of ∼10^4^ when cells were grown on PET (**Figure 2B&D** and **Figure S9B&D**). Peaks for MHET were detected above the baseline (*i.e.,* ∼ 10^5^ vs a baseline of ∼10^4^), however the areas were too low to quantify using our calibration curve (**Figure 2C** and **Figure S9C**). Despite not being able to quantify MHET, we observed a temporal trend where the highest peak areas occurred between weeks 2 and 4, which coincided with maximum cell growth for each culture (**Figure S10**). These observations suggest MHET was being produced at low levels and that cells likely converted this MHET into other metabolites to support growth thereby precluding quantification.

**Figure 2:**
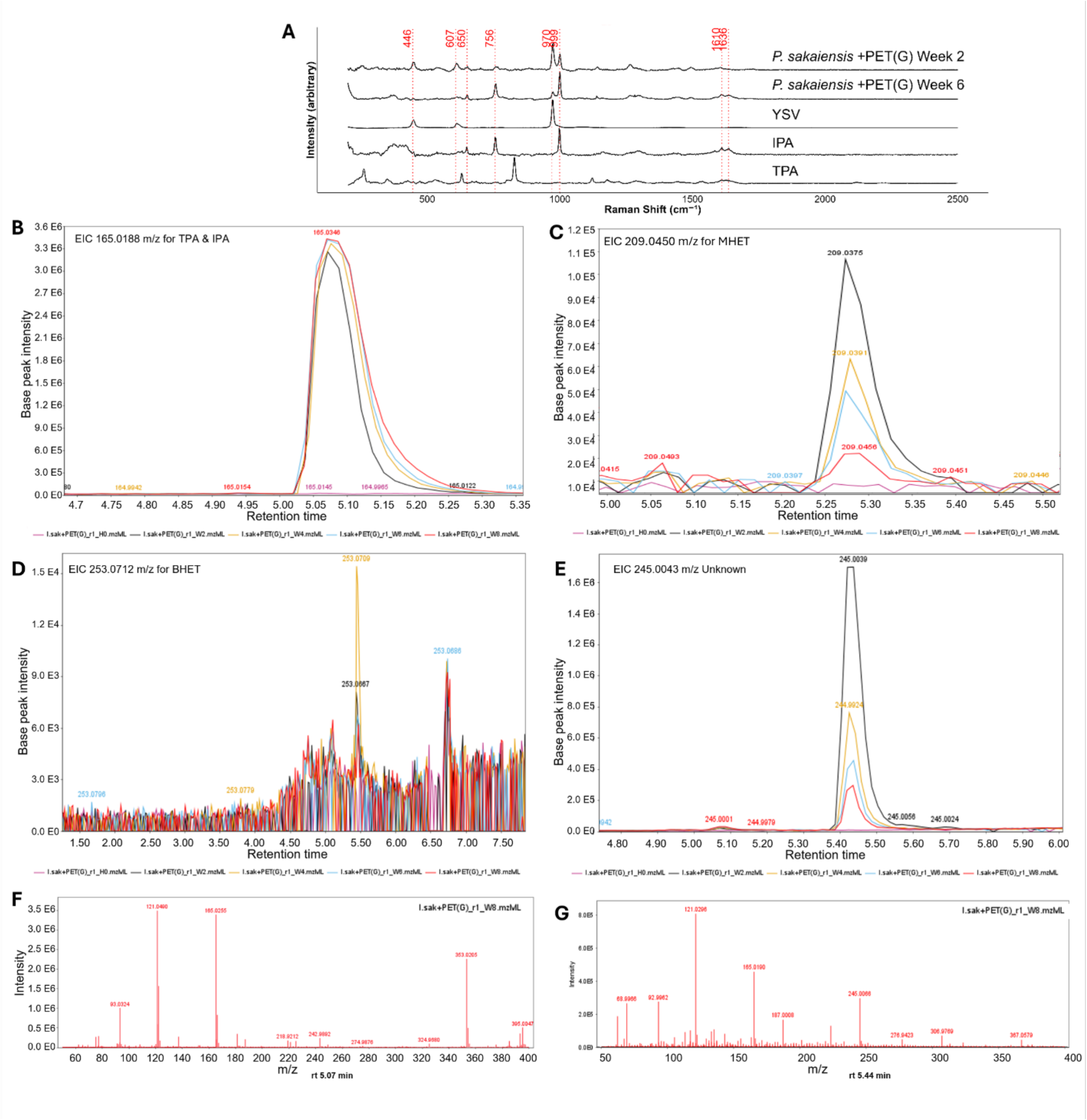
Detection of *P. sakaiensis* monomers from the metabolism of polyethylene terephthalate (PET) manufactured by Goodfellows (G) during the 8-week incubation. (A) Raman spectroscopy data comparison for cell culture samples at week 2 and week 6 and isophthalic acid (IPA) and terephthalic acid (TPA) standards. (B-E) are the extracted ion chromatograms (EIC) of samples obtained from cultures from week 0 to week 8 for *m/z* 165.0188 (B), 209.0450 (C), 253.0712 (D) and 245.0043 for the unknown compound (E). (F) and (G) are the mass spectra (with in-source fragmentation) for IPA at RT of 5.07 min (G) and the unknown compound at RT of 5.44 min (G). EIC were constructed using *m/z* tolerance of 0.01.

Although we expected MHET to be converted partially to TPA based on the current understanding of PET metabolism in this model organism, we did not detect any TPA in our cultures. Instead, we consistently detected one metabolite sharing the same *m/z* value that eluted at a later RT than that of TPA (*i.e.*, 5.07 min vs 4.75 min) [**Figure 2B** for PET(G) and **Figure S9B** for PET(M)]. We suspected the shift in RT for compounds sharing the same *m/z* value was because these compounds were isomers. Using the standards for benzene dicarboxylic acid isomers (*i.e.*, TPA and IPA), we confirmed the metabolite with an *m/z* of 165.0188 was IPA (**Figure S1**). The concentration of IPA increased from 0 to ∼11 ppm and 0 to ∼58 ppm at week 4 when cells were grown on PET(M) and PET(G), respectively, and reached ∼15 and 73 ppm towards the end of the incubation period of 8 weeks (**Figure 3**).

**Figure 3:**
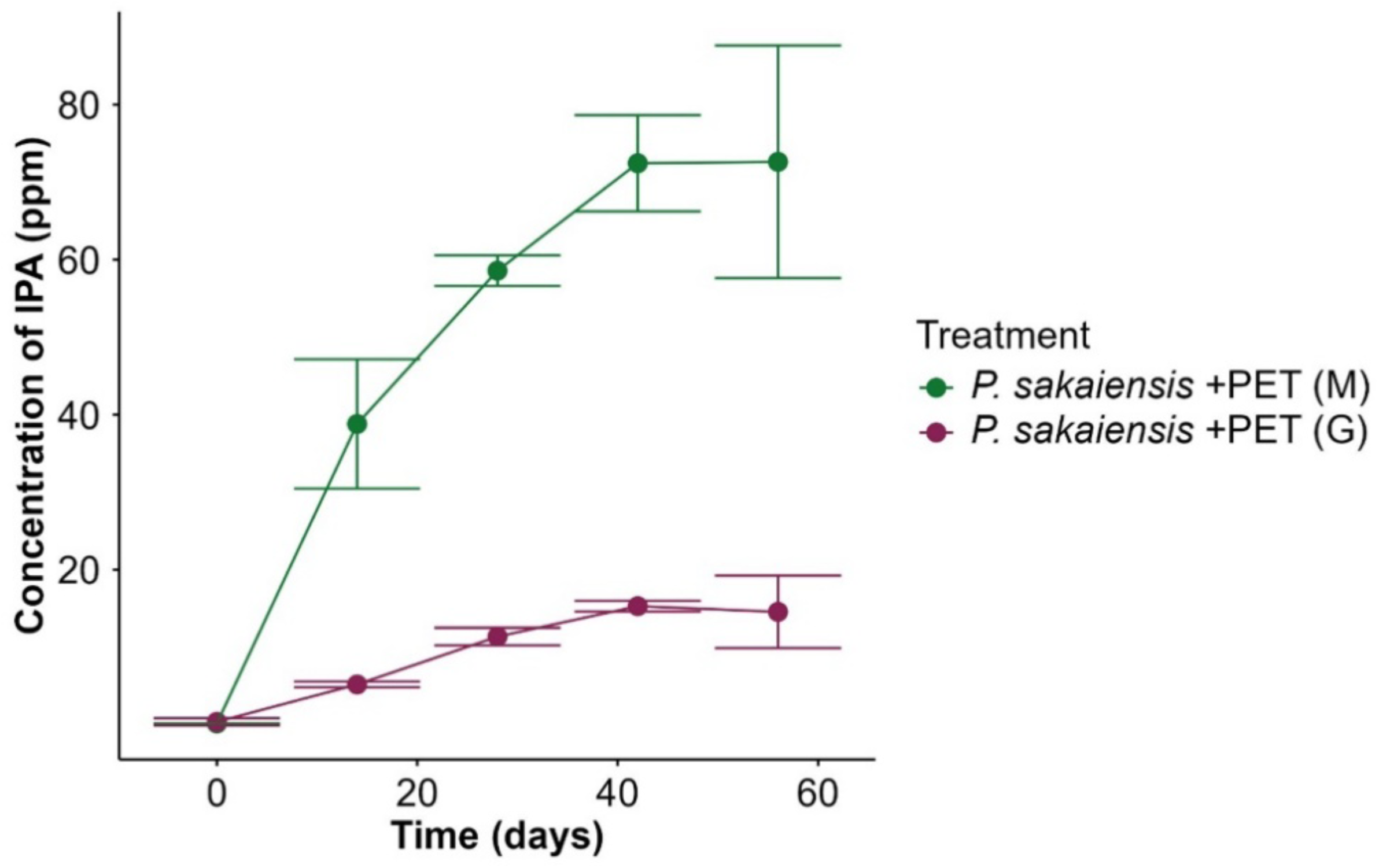
Detection of isophthalic acid (IPA) (ppm ± SD, n=3) from *P. sakaiensis* grown on ground Goodfellow (G) and Magerial Science (M) sourced polyethylene terephthalates (PET).

PET metabolites including IPA were also analyzed using Raman spectroscopy. TPA and its precursors MHET and BHET share many similar band assignments, with additional peaks occurring in MHET and BHET due to their more complex molecular structure. Although IPA and TPA have the same chemical formula, their functional group arrangements produce different Raman spectra (**Figure S4**) with different band assignments bearing no overlap (**Table S5** and **Table S6**). The feature bands for TPA, MHET and BHET were not observed in the PET biodegradation experiment while Raman bands matching IPA were detected (**Figure 2A** and **Figure S9A**). We suspect the lack of detection of MHET via Raman is due to the inherently lower sensitivity of this method, which is inefficient at detecting chemicals <1 ppm (Hu et al., 2015).

At week 2, the Raman spectra for live cells supplied with PET(G) exhibited the 3 culture medium associated peaks, with 970 cm^-1^ being the strongest peak along with IPA related peaks at 756 and 999 cm^-1^ (**Figure 2A**). When IPA production plateaued at week 6 (**Figure 3**), the PET(G) spectra had more IPA related peaks at 650, 756, 1610 and 1636 cm^-1^, with 999 cm^-1^ becoming the strongest peak rather than 970 cm^-1^ (**Figure 2A**). In contrast, the peak at 999 cm^-1^ for samples obtained from cultures supplied with PET(M) remained consistently low and did not surpass the intensity of the peak at 970 cm^-1^ (**Figure S9A**). The latter observation aligns with the lower IPA concentrations measured via LC-HRMS when cells were grown on PET(M) (**Figure 3**).

In contrast to IPA, no EG was detected in the supernatants of cell cultures grown on PET using GC-MS or Raman spectroscopy. Experiments where EG was supplied as a carbon source showed that EG concentrations in culture supernatants decreased from ∼20 to 3 ppm from day 0 to 14 (**Figure S11A&C**). These observations align with cells taking up EG to support modest growth observed early in the incubation as seen in other studies (Kalathil et al., 2022) (**Figure 1B** and **Figure S11**). Experiments carried out with TPA showed minimal change in the TPA signal suggesting no TPA was being taken up or metabolized by the cells (**Figure S11 B&D**).

In addition to IPA, LC-HRMS analyses revealed another peak corresponding to an uncharacterized metabolite that eluted at 5.43 min with a *m/z* of 245.0039 *m/z* (**Figure 2E&G** and **Figure S9E&G**). The peak area of this compound decreased after week 2 when cells were provided PET(G) but increased in the first 4 weeks when cells were provided PET(M) followed by a decrease at the end of the cultivation period (**Figure 2E** and **Figure S9E**). The in-source fragments or adducts of this unknown compound were similar to those observed for IPA (*i.e.*, 121.0371 *m/z* and 165.0367 *m/z*) with the addition of a unique fragment detected at 223.0331 *m/z*. Although the RT of this compound was similar to BHET (**Figure S1A**), the in-source fragmentation pattern did not match (**Figure 2G** and **Figure S9G**).

The similarities between this unknown metabolite and IPA suggest the compound might be a precursor of IPA. The accumulation and subsequent depletion of this putative intermediate coincided with increases in IPA concentration (**Figure S12**), suggesting metabolic conversion. The timing in the accrual of this intermediate compared to IPA supports this explanation. Cells provided with PET(G) exhibited rapid intermediate formation and earlier IPA accumulation compared to cells provided PET(M) (**Figure 2E, Figure S9E** and **Figure 3**). This delay may reflect differences in polymer physico-chemical properties (*e.g*., surface area and crystallinity) that affect the capacity for cells to access PET and initiate metabolism.

### PET crystallinity measurements via Raman spectroscopy

Determining whether polymers have low crystallinity and high bioavailability vs high crystallinity and low bioavailability is important for predicting plastic biodegradation (Cai et al., 2023; Kong & Hay, 2002; Tokiwa et al., 2009). To assess Raman’s capacity to link crystallinity to biodegradability, we collected Raman spectral information for four different PET materials: The two sources of PET used in our cell growth experiments (ground Goodfellow and Magerial Science) and an additional Goodfellow amorphous film and Goodfellow biaxially orientated film (**Methods**). We independently measured crystallinity for these materials using DSC to compare analytical methods and provide more context for the metabolite analyses summarized previously.

The crystallinity values obtained by DSC were 43.41% for Magerial Science PET powder (**Figure S13A**), 33.00% for Goodfellow biaxially orientated film (**Figure S13B**), 7.44% for Goodfellow amorphous film ground (**Figure S13C**) and 7.79% for Goodfellow amorphous film (**Figure S13D**). Raman spectra for all four types of material contained similar band assignments typically associated with PET (table inset to **Figure 4**) with variations in peak intensity for the band assignments used to measure crystallinity (spectra inset in **Figure 4**). The lower crystallinity PET film and ground amorphous PET from Goodfellow had weaker signals at 1092 cm^-1^ compared to the high crystallinity materials (**Figure 4**), resulting in a lower 1092/1114 cm^-1^ ratio. For example, the ratio for amorphous PET from Goodfellow was ∼0.6 compared to the higher crystallinity PET from Magerial Science, which was ∼1.7 (**Table S7**). Low ratios between these two band assignments correspond to lower crystallinity showing good alignment between our Raman and DSC analyses (Galea et al., 2025; Lin et al., 2016).

**Figure 4:**
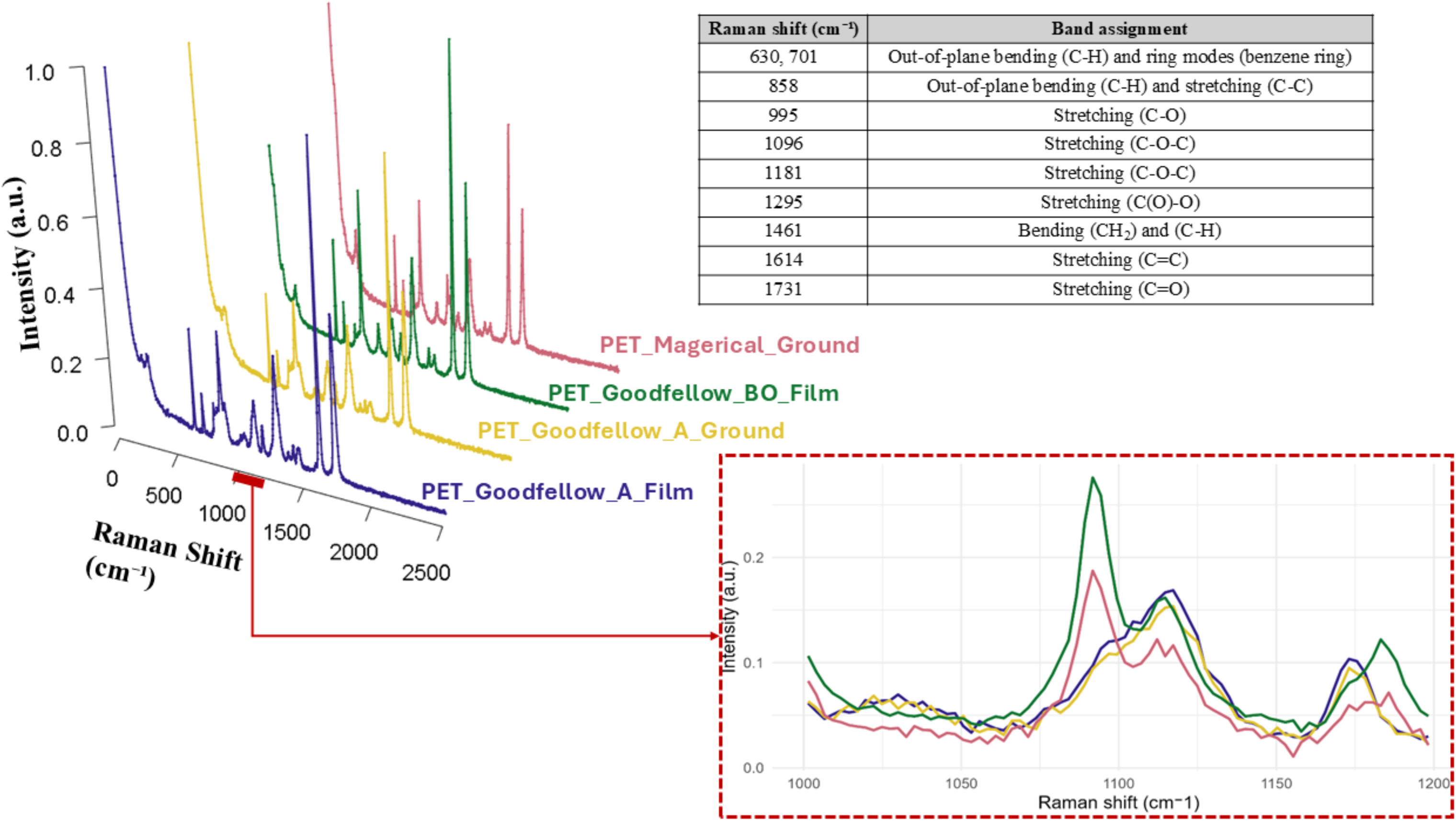
Raman spectrum of various manufacturers and orientations of polyethylene terephthalate (PET) used for PET degradation experiments with zoom on 1092 and 1114 cm^-1^ peaks. The materials included are from Magerial Science manufacturer ground PET powder (PET_Magerial_Ground), Goodfellow manufacturer bilateral orientation PET film (PET_Goodfellow_BO_Film), Goodfellow manufacturer amorphous PET film which was ground (PET_Goodfellow_A_Ground) and Goodfellow manufacturer amorphous PET film (PET_Goodfellow_A_Film). The table is the band assignments of PET with the corresponding Raman shifts (Peñalver et al., 2023).

These experiments provided insights into how increasing the surface area of PET materials with similar crystallinity can affect biodegradation. When comparing the ground and unmanipulated films from Goodfellow that had similar low crystallinities (*i.e.,* ∼ 7 to 8%) the ground material led to detectable metabolite production whereas the film did not (**Figure 2** and **Figure S14**). We also observed that PET biodegradability and IPA production were inversely related to crystallinity, similar to previous work (Di Lorenzo, 2024). The ground amorphous PET film from Goodfellow had considerably lower crystallinity (*i.e.,* 7.44 % for PET(G)) than the Magerial Science powder (*i.e.,* 43.41 % for PET(M)) and resulted in higher IPA production over time (*i.e.,* ∼73 ppm vs ∼15 ppm) (**Figure 3**). Notably, there was still evidence that cells grown on PET(M) released IPA despite its considerably higher crystallinity (**Figure 3**). We suspect cells maintained PET metabolism because the powder form of PET(M) increased surface area, which can overcome barriers PET hydrolases encounter when transforming highly crystalline substrates (Erickson et al., 2022).

## Discussion

This study demonstrates that Raman spectroscopy can provide molecular and physical structural data for PET alongside rates of metabolite production indicative of plastic biodegradation. This study shows how Raman complements gold standard analytical techniques to provide novel mechanistic insights into even well-studied plastic metabolisms. One of the more impactful findings in this study was the detection of IPA via independent analytical techniques. The detection of IPA occurred alongside deviations from the expected physiological role of TPA. Based on this finding, we think it is valuable to reexamine previously published work and reconsider the mechanistic understanding of PET metabolism in this model organism.

Multiple studies have detected TPA as a product of PET metabolism, although these observations are primarily from work on isolated enzymes (Moog et al., 2019; Kim et al., 2020; Chen et al., 2022; Wang et al., 2022; Boros et al., 2025). Within enzymatic studies, there is evidence that TPA and EG can inhibit the initial reactions catalyzed by PETase (Erickson et al., 2022). These observations contrast with reports using live cells where TPA and EG were used as substrates for growth and to stimulate PET metabolism (Hachisuka et al., 2022; Poulsen & Nielsen, 2023; Tanaka et al., 2024; Zhao et al., 2025).

In the original study where TPA was characterized as an intermediate of PET metabolism, the authors did not detect TPA based on the standards used in their HPLC-UV analyses (Yoshida et al., 2016). Instead, the authors posit that TPA could not be detected due to *P. sakaiensis* metabolizing it. In another study, TPA could not be detected in culture supernatants with *P. sakaiensis* grown anaerobically on PET after a 30 day incubation using NMR spectroscopy (Kalathil et al., 2022). In the latter case, intracellular metabolism of TPA could not be eliminated as a possibility, but this seems unlikely in our experiments because there was no evidence of TPA uptake. Our observations of TPA inhibiting cell growth are better aligned with a short-term (*i.e.,* ∼ 24 hours) proteomic study which suggested TPA can inhibit chaperone proteins and central carbon metabolism in *P. sakaiensis* (Poulsen & Nielsen, 2023). Whether this specific mode of action contributes to TPA toxicity needs to be formally tested.

The comparisons cited above show there are still facets of this model organism’s PET metabolism that need to be resolved. Although some of the discrepancies likely arise from differences in the biological systems and experimental setups employed, these inconsistencies persist in studies that adhere to the conditions in the seminal work on *P. sakaiensis* (Hachisuka et al., 2022; Piedra et al., 2025; Poulsen and Nielsen, 2023; Tanasupawat et al., 2016; Wallace et al., 2020; Walter et al., 2022; Yoshida et al., 2016). Given these differences, it is valuable to consider the physiological context in which IPA could be produced because of PET metabolism.

Producing IPA as a metabolic intermediate would provide cells with a more bioavailable substrate compared to TPA given IPA’s higher water solubility (*i.e.*, 120 mg/L for IPA vs 17 mg/L for TPA at 25°C) (Park & Sheehan, 2000). Our data indicate that IPA concentrations plateau shortly after cells reached peak density, suggesting IPA production is tied to cellular metabolism (**Figure 1A** and **Figure 3**). Our data also suggest there is an unidentified precursor to IPA that is likely an iso-form of MHET that decreases as IPA is formed (**Figure S13**), which is indicative of potential metabolic connections that merit further investigation.

We acknowledge that IPA could also be released because it is used to produce TPA (2 wt %) (Abedsoltan, 2023) and it is incorporated into PET to decrease crystallinity (Tomisawa et al., 2022; Roy & Pellerin, 2025). As such, IPA may simply be released as the crystalline PET matrix is depolymerized rather than a metabolic reaction that produces IPA. This explanation is plausible based on chemical PET transformation experiments where hydrolysis with alkaline and acid treatments released benzoic acid, BHET, IPA, and TPA in the aqueous phase (Thulasiraman et al., 2025). Our growth experiments indicate that cells accessed similar amounts of carbon on low and high crystallinity PET (**Figure 1A**). As such, the differences in IPA produced on PET(G) vs PET(M) could be a byproduct of different amounts of IPA being present in low and high crystallinity PET materials (**Figure 3**). Confirming if IPA is a genuine metabolite produced during PET metabolism or released due to depolymerization will be key to test in future experiments.

## Conclusion

This study frames Raman spectroscopy as a powerful tool that provides real-time monitoring data on the physical and biological controls affecting plastic biodegradation. We show that Raman spectroscopy can be used to rapidly screen cultures for known and potentially novel plastic biodegradation metabolites (*e.g.,* IPA). This screening capability is valuable for guiding subsequent analytical efforts to provide additional molecular structure details using mass spectrometry. These complimentary approaches could advance our mechanistic understanding of an ever-growing catalogue of plastic degrading microbes. Carrying out such fundamental studies would lay the groundwork for using Raman spectroscopy as a PAT to characterize how physicochemical properties of different plastics (*e.g.,* crystallinity, polymer identity, presence of additives) affect biodegradation. Leveraging these insights as part of quantitative models to predict the biodegradability of mixed plastic streams will be critical to enhancing the overall performance of biological plastic reclamation *in situ*. In that respect, we consider Raman spectroscopy as a key tool that can support a more circular plastic economy.

## Associated Content

**Supporting Information 1**. Additional experimental details and results including tables and figures (Word).

## Acknowledgments

SNP contributed to the conceptualization, methodology, investigation, formal analysis, validation, writing the original draft. YH contributed to the conceptualization, data curation, formal analysis, project administration, resources, software, supervision, LC-HRMS methodology and Raman methodology and instrumentation. DRM contributed to the LC-HRMS methodology and instrumentation. PH contributed to the LC-HRMS methodology and investigation. AB contributed to the GC-MS methodology. DSG contributed to the conceptualization, data curation, formal analysis, project administration, resources, software, and supervision. All authors contributed to reviewing and editing the manuscript. This work was supported by an Environmental Research and Education Foundation Graduate scholarship to SNP, an NSERC Discovery Grant awarded to DSG (#RGPIN-2022-04891 and DGECR-2022-00512), NSERC Discovery Grant and RTI awarded to YH (#RGPIN-2022-04892 and RTI-2024-00300) and NSERC Alliance (ALLRP 602597-25) awarded to DSG and YH. We would like to thank Dr. Banu Ormeci (Department of Engineering, Carleton University) for providing us access to their GC-MS instrument.

## Abbreviations

BHET: bis-(2-hydroxyethyl) terephthalate
DSC: differential scanning calorimeter
EG: ethylene glycol
GC-MS: gas chromatography-mass spectroscopy
IPA: isophthalic acid
LC-HRMS: liquid chromatography-high resolution mass spectroscopy
MHET: mono-(2-hydroxyethyl) terephthalate
PAT: process analytical technologies
PET: polyethylene terephthalate
RT: retention times
TPA: terephthalic acid.

---

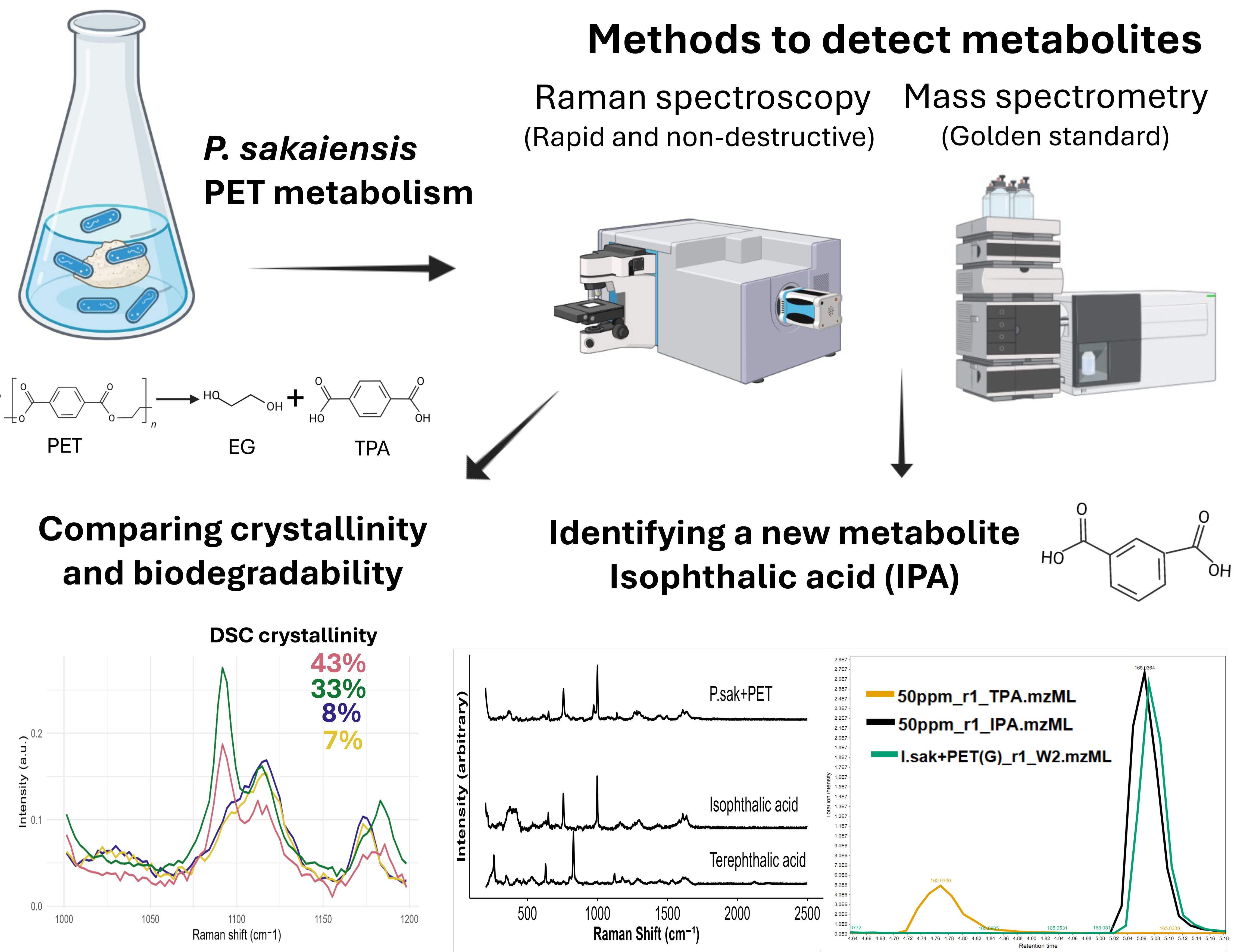

