## Supplementary Information for "Raman Spectroscopy Enables Real-Time Identification and Monitoring of Plastic Biodegradation Metabolites"

**Figure S7**: Extracted Ion Chromatograph of abiotic controls throughout the 8-week degradation experiment.................................................................................................................................. 12

**Figure S8**: Raman spectrums of controls for degradation experiment at week 2. ....................... 13

**Figure S9**: Detection of *P. sakaiensis* degradation of Magerial manufactured PET monomers. . 14

**Figure S10**: Detection of the *P. sakaiensis* monomer MHET during cellular growth from the metabolism of group up PET(G)……………………...…………………………………………15

**Table S5.** Terephthalic acid Raman spectrum band assignments ............................................. 166

**Table S6**. Isophthalic acid Raman spectrum band assignments ............................................... 166

**Figure S11**: Extracted Ion Chromatograph of *P. sakaiensis* degradation of various monomers ..............................................................................................**Error! Bookmark not defined.**7

**Figure S12:** Detection of the *P. sakaiensis* monomer IPA compared to an unknown metabolite during cellular growth from the metabolism of group up PET(G)…………………..………........... 18

**References**……………………………..………………………………………………..………21

**Medium recipes**

Reasoner's 2A (R2A) medium: 0.5 g $L^{-1}$ yeast extract (Bioshop), 0.5 g $L^{-1}$ proteose peptone (Gibco), 0.5 g $L^{-1}$ casamino acid (Thermo Scientific), 0.5 g $L^{-1}$ glucose (Bioshop), 0.5 g $L^{-1}$ starch (Thermo Scientific), 0.3 g $L^{-1}$ Na-pyruvate (Sigma-Aldrich), 0.3 g $L^{-1}$ potassium phosphate dibasic [$K_2HPO_4$] (Sigma-Aldrich), 0.05 g $L^{-1}$ magnesium sulfate heptahydrate [$MgSO_4 \cdot 7H_2O$] (Sigma-Aldrich), and 15 g $L^{-1}$ agar (Bioshop) in 1 L of milliQ water with a pH of 7.2 using sodium phosphate dibasic [$Na_2HPO_4$] or sodium phosphate monobasic [$NaH_2PO_4$] to adjust the pH.

Yeast extract-sodium carbonate-vitamins (YSV) medium: 0.1 g $L^{-1}$ yeast extract (Bioshop), 0.2 g $L^{-1}$ sodium bicarbonate (Sigma-Aldrich), 1 g $L^{-1}$ ammonium sulfate (Thermo Scientific), 0.1 g $L^{-1}$ calcium carbonate (Bioshop), 1 L of milliQ water with a pH of 7.0], plus a trace element mix [1 g $L^{-1}$ iron (III) sulfate heptahydrate [$FeSO_4 \cdot 7\ H_2O$] (Thermo Scientific), 0.1 g $L^{-1}$ copper(II) sulfate pentahydrate [$CuSO_4 \cdot 5\ H_2O$] (Thermo Scientific), 0.1 g $L^{-1}$ zinc sulfate heptahydrate [$ZnSO_4 \cdot 7\ H_2O$] (Thermo Scientific), 0.61 g $L^{-1}$ magnesium sulfate hydrate [$MnSO_4 \cdot H_2O$] (Sigma-Aldrich)] and a vitamin mix [2.5 g $L^{-1}$ Thiamine Hydrochloride (Sigma-Aldrich), 0.05 g $L^{-1}$ biotin (TCI), 0.5 g $L^{-1}$ vitamin B12 (Thermo Scientific)] in 10 mM phosphate buffer (sodium phosphate dibasic heptahydrate [$Na_2HPO_4 \cdot 7\ H_2O$] (Sigma-Aldrich) and sodium phosphate monobasic monohydrate [$Na2H_2PO4 \cdot H_2O$] (Sigma-Aldrich)

**Table S1**. Experimental setup of polyethylene terephthalate (PET) and monomers (terephthalic acid (TPA) and ethylene glycol (EG)) degradation (n=3).

| Name | Contents | Group Type |
|---|---|---|
| YSV | YSV medium only | Medium control |
| YSV + EG | YSV + EG | Abiotic control |
| YSV + TPA | YSV + TPA | Abiotic control |
| YSV + PET-M | YSV + PET (Material Science) | Abiotic control |
| YSV + PET-G | YSV + PET (Goodfellow) | Abiotic control |
| *P. sakaiensis* | YSV medium + washed bacterial culture | Biotic control |
| *P. sakaiensis* + glucose | YSV medium + glucose + washed bacterial culture | Alternate carbon source control |
| *P. sakaiensis* + EG | YSV medium + EG + washed bacterial culture | Sample |
| *P. sakaiensis* + TPA | YSV medium + TPA + washed bacterial culture | Sample |
| *P. sakaiensis* + PET-M | YSV medium + PET-M + washed bacterial culture | Sample |
| *P. sakaiensis* + PET-G | YSV medium + PET-G+ washed bacterial culture | Sample |

**Table S2**. Reverse phase liquid-chromatography high resolution mass spectrometry binary pump timetable. (A) ddH2O with 0.1% formic acid and (B) acetonitrile (ACN) with 0.1 % formic acid.

| Time (min) | A (%) | B (%) |
|---|---|---|
| 0.00 | 95.0 | 5.0 |
| 1.00 | 95.0 | 5.0 |
| 2.00 | 70.0 | 30.0 |
| 5.00 | 50.0 | 50.0 |
| 8.00 | 0.0 | 100.0 |
| 9.00 | 95.0 | 5.0 |
| 10.00 | 95.0 | 5.0 |

**Table S3**. Summary of breakdown products of polyethylene terephthalate (PET) standards including bis-(2-hydroxyethyl) terephthalate (BHET), mono-(2-hydroxyethyl) terephthalate (MHET), terephthalic acid (TPA), isophthalic acid (IPA) and ethylene glycol (EG) with mass-to-charge ratios (*m/z*) and retention times ran on either the LC-HRMS and GC-MS. While BHET, MHET, TPA and IPA are deprotonated molecule ($[M\text{-}H]^{-1}$), EG is the molecular ion ($[M]^{+}$),

| Metabolite | Molecular Formula | Mass-to-charge ratio (*m/z*) | Retention time (minutes) |
|---|---|---|---|
| BHET | C12H14O6 | 253.0712 | 5.44 |
| MHET | C10H10O5 | 209.0450 | 5.24 |
| TPA | C8H6O4 | 165.0188 | 4.77 |
| IPA | C8H6O4 | 165.0188 | 5.07 |
| EG | C2H6O2 | 61.0290 | 5.84 |

A

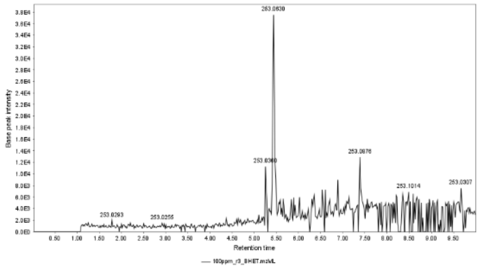

B

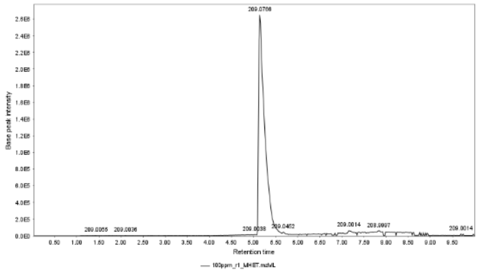

C

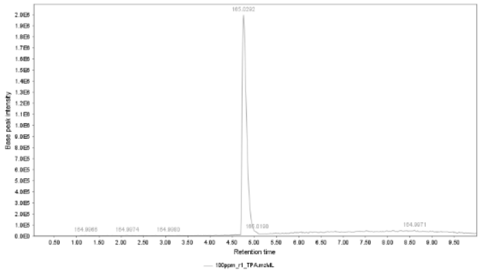

D

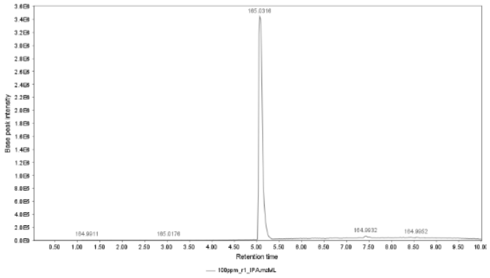

E

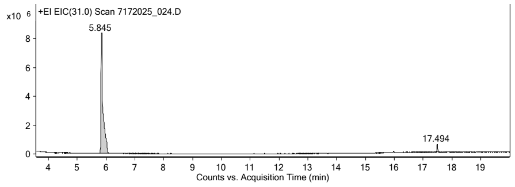

**Figure S1**: Extracted Ion Chromatograph (EIC) of all breakdown products of polyethylene terephthalate (PET) standards at 100 ppm including (A) bis-(2-hydroxyethyl) terephthalate (BHET), (B) mono-(2-hydroxyethyl) terephthalate (MHET), (C) terephthalic acid (TPA), (D) isophthalic acid (IPA) and (E) ethylene glycol (EG).

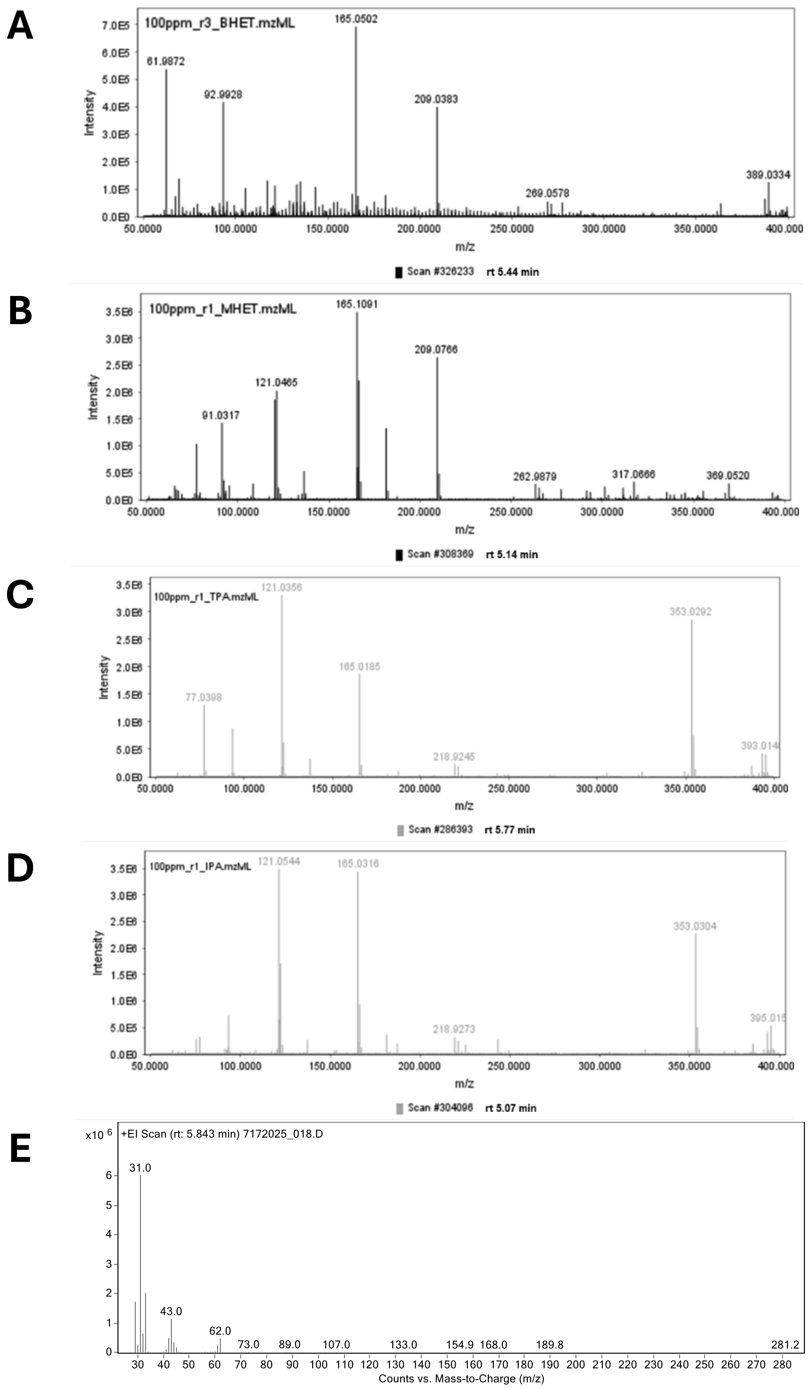

**Figure S2**: Mass spectrum of expected breakdown products of polyethylene terephthalate (PET) standards including (A) bis-(2-hydroxyethyl) terephthalate (BHET), (B) mono-(2-hydroxyethyl) terephthalate (MHET), (C) terephthalic acid (TPA), (D) isophthalic acid (IPA) and (E) ethylene glycol (EG).

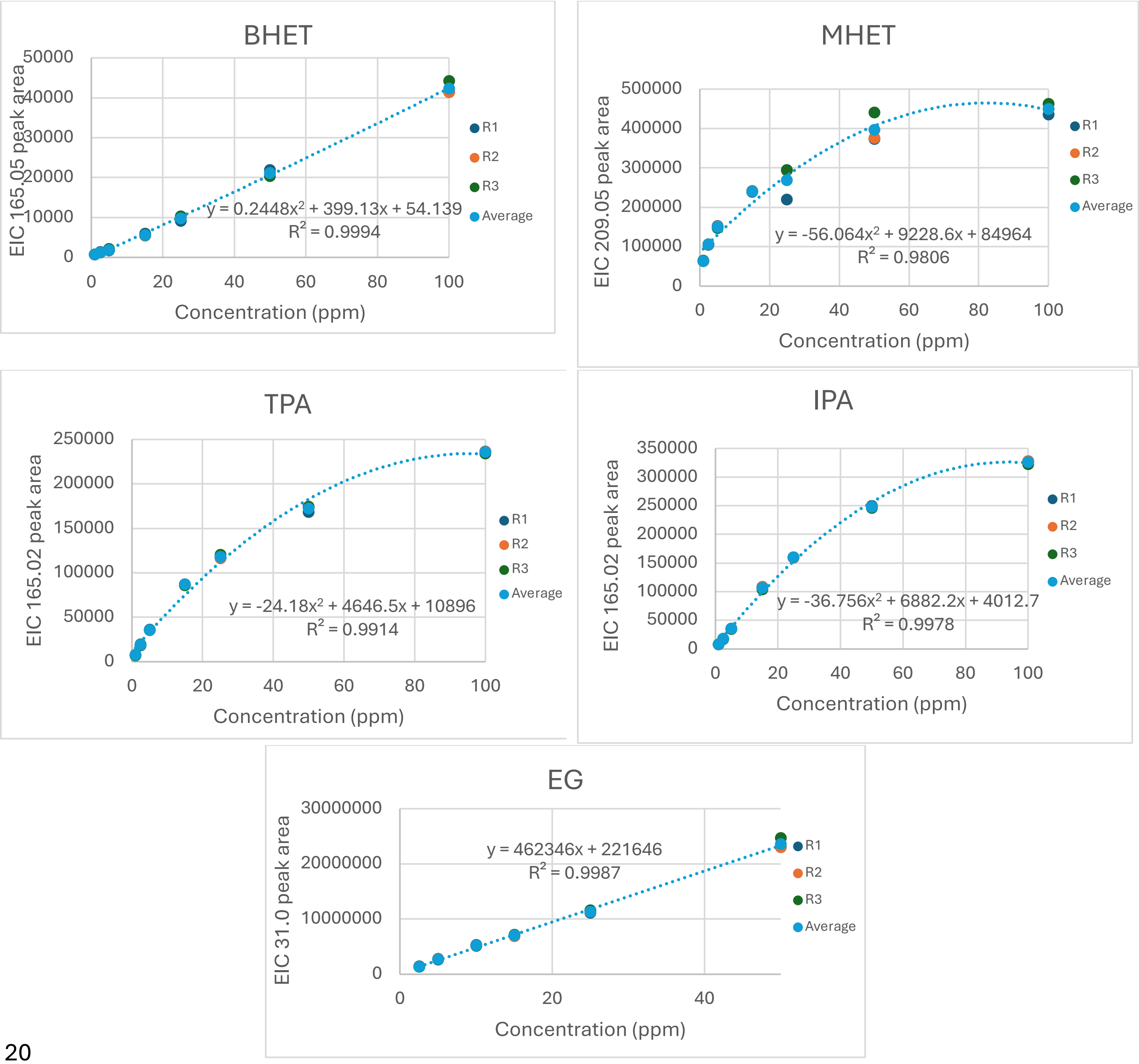

**Figure S3**: Calibration curves with replicates (n = 3) and averages of breakdown products of polyethylene terephthalate (PET) standards ran on either the LC-HRMS and GC-MS including bis-(2-hydroxyethyl) terephthalate (BHET), mono-(2-hydroxyethyl) terephthalate (MHET), terephthalic acid (TPA), isophthalic acid (IPA) and ethylene glycol (EG) using quadradic or linear trendlines.

**Table S4**. Regression equations for standard curves for breakdown products of polyethylene terephthalate (PET) standards ran on LC-HRMS and GC-MS including bis-(2-hydroxyethyl) terephthalate (BHET), mono-(2-hydroxyethyl) terephthalate (MHET), terephthalic acid (TPA), isophthalic acid (IPA) and ethylene glycol (EG). Standards run on the LC-HRMS ranged from 1 to 100 mg/L. Standards run on the GC-MS ranged from 2.5 to 100 mg/L.

| **Metabolite** | **Regression line (mg/L)** | **Regression coefficient ($R^2$)** |
|---|---|---|
| BHET | $0.2448x^2 + 399.13x + 54.139$ | 0.9994 |
| MHET | $-56.064x^2 + 9228.6x + 84964$ | 0.9806 |
| TPA | $-24.18x^2 + 4646.5x + 10896$ | 0.9914 |
| IPA | $-36.756x^2 + 6882.2x + 4012.7$ | 0.9978 |
| EG | $462346x + 221646$ | 0.9987 |

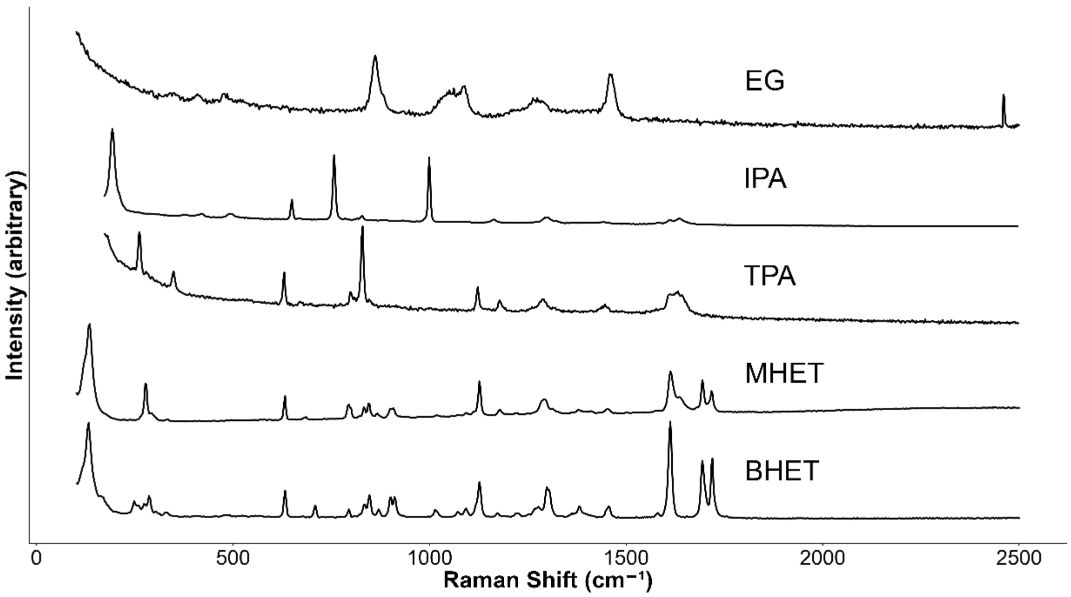

**Figure S4**: Raman spectra of standards for breakdown products of polyethylene terephthalate (PET) including bis-(2-hydroxyethyl) terephthalate (BHET), mono-(2-hydroxyethyl) terephthalate (MHET), terephthalic acid (TPA), isophthalic acid (IPA) and ethylene glycol (EG).

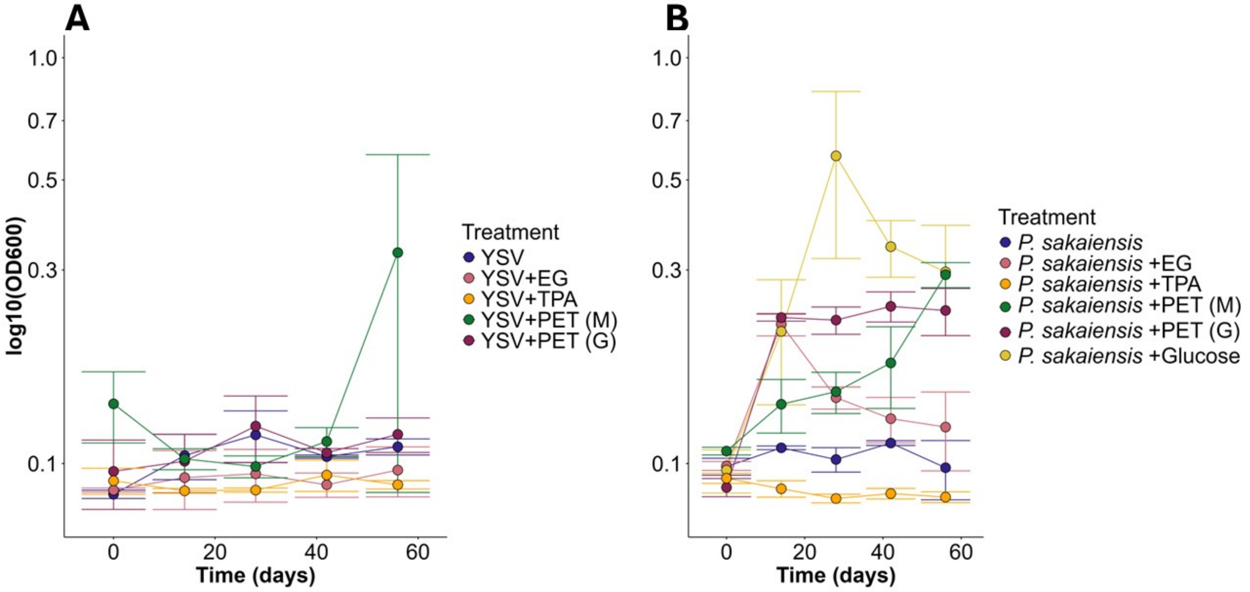

**Figure S5**: Cell density of *P. sakaiensis* measured using the log10 of $OD_{600}$ (± SD, N=3) throughout the 8-week degradation experiments (A) without and (B) with cells. Various treatments including YSV medium, ethylene glycol (EG), terephthalic acid (TPA), polyethylene terephthalate (PET) manufactured by Magerial (M), polyethylene terephthalate (PET) manufactured by Goodfellows (G) and glucose.

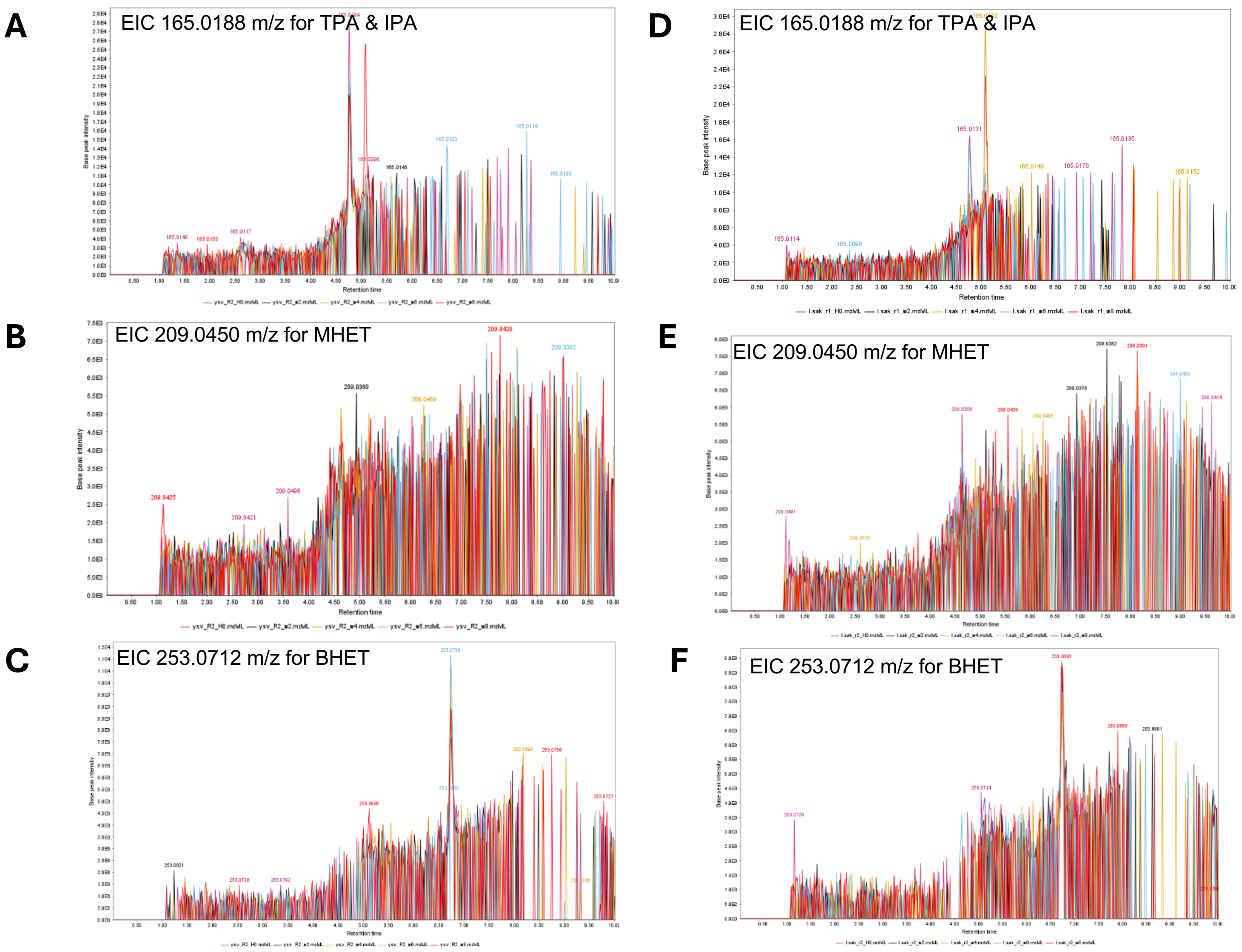

**Figure S6**: Representative extracted ion chromatograph (EIC) of LC-MS results for biological controls for degradation experiment at weeks 0, 2, 4, 6, and 8. (A-C) are for YSV medium and (D-F) are biotic (*P. sakaiensis*) in YSV medium. EICs were constructed with *m/z* tolerance of 0.01 Da for *m/z* values of 165.0188 (A and D); 209.0450 (B and E); and 253.0712 (C and F), corresponding to terephthalic acid (TPA) and isophthalic acid (IPA); monohydroxyethyl terephthalate (MHET); and Bis(2-hydroxyethyl) terephthalate (BHET), respectively.

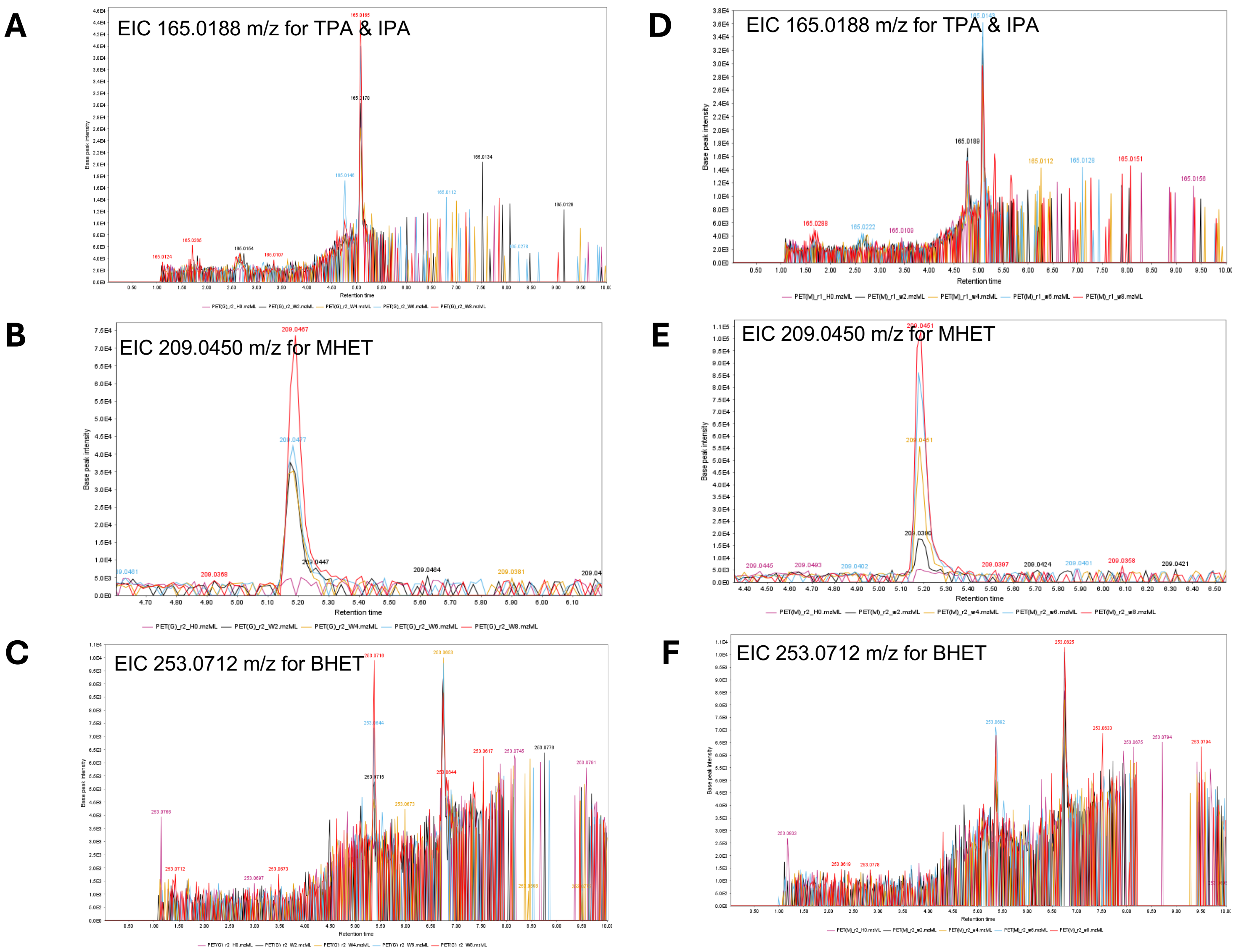

**Figure S7**: Representative extracted ion chromatograph (EIC) of LC-MS results for abiotic plastic controls for degradation experiment at weeks 0, 2, 4, 6, and 8. (A-C) are for polyethylene terephthalate (PET) manufactured by Goodfellows (G) and (D-E) are for PET manufactured by Magerial Science (M). EICs were constructed with *m/z* tolerance of 0.01 Da for *m/z* values of 165.0188 (A and D); 209.0450 (B and E); and 253.0712 (C and F), corresponding to terephthalic acid (TPA) and isophthalic acid (IPA); monohydroxyethyl terephthalate (MHET); and Bis(2-hydroxyethyl) terephthalate (BHET), respectively.

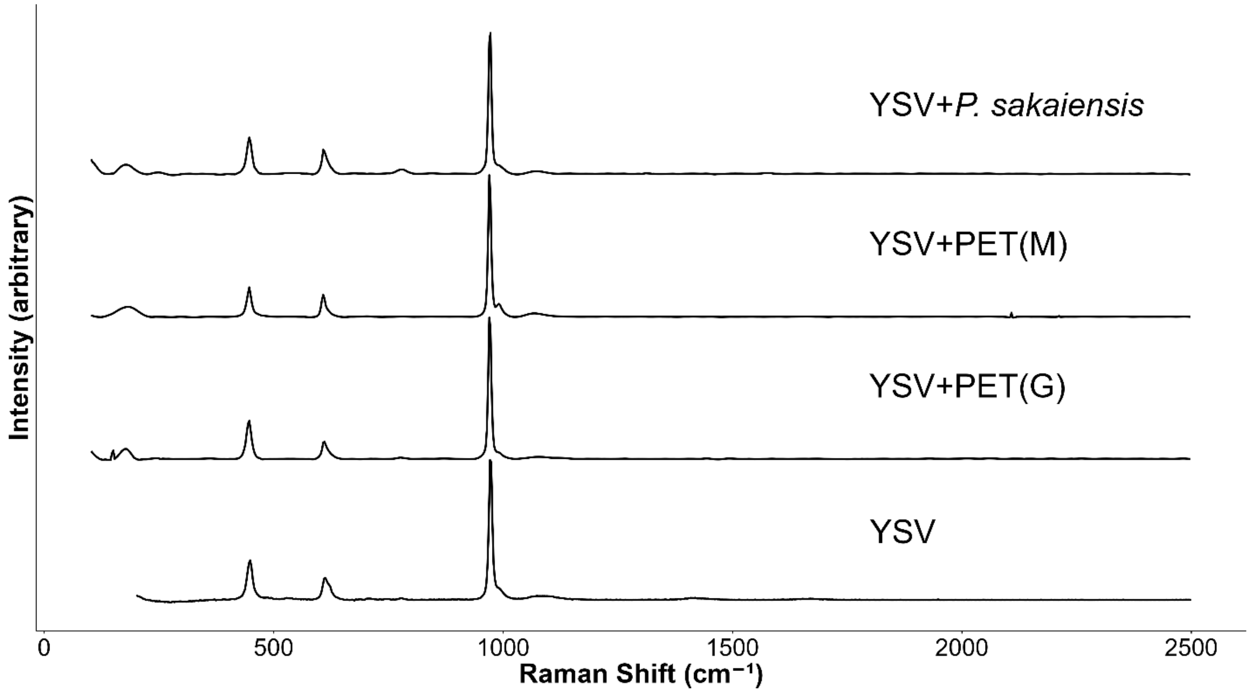

**Figure S8**: Raman spectra of controls for degradation experiment at week 2 including medium (YSV medium), abiotic (polyethylene terephthalate (PET) manufactured by Goodfellows (G) and Magerial Science (M)) and biotic (*P. sakaiensis*).

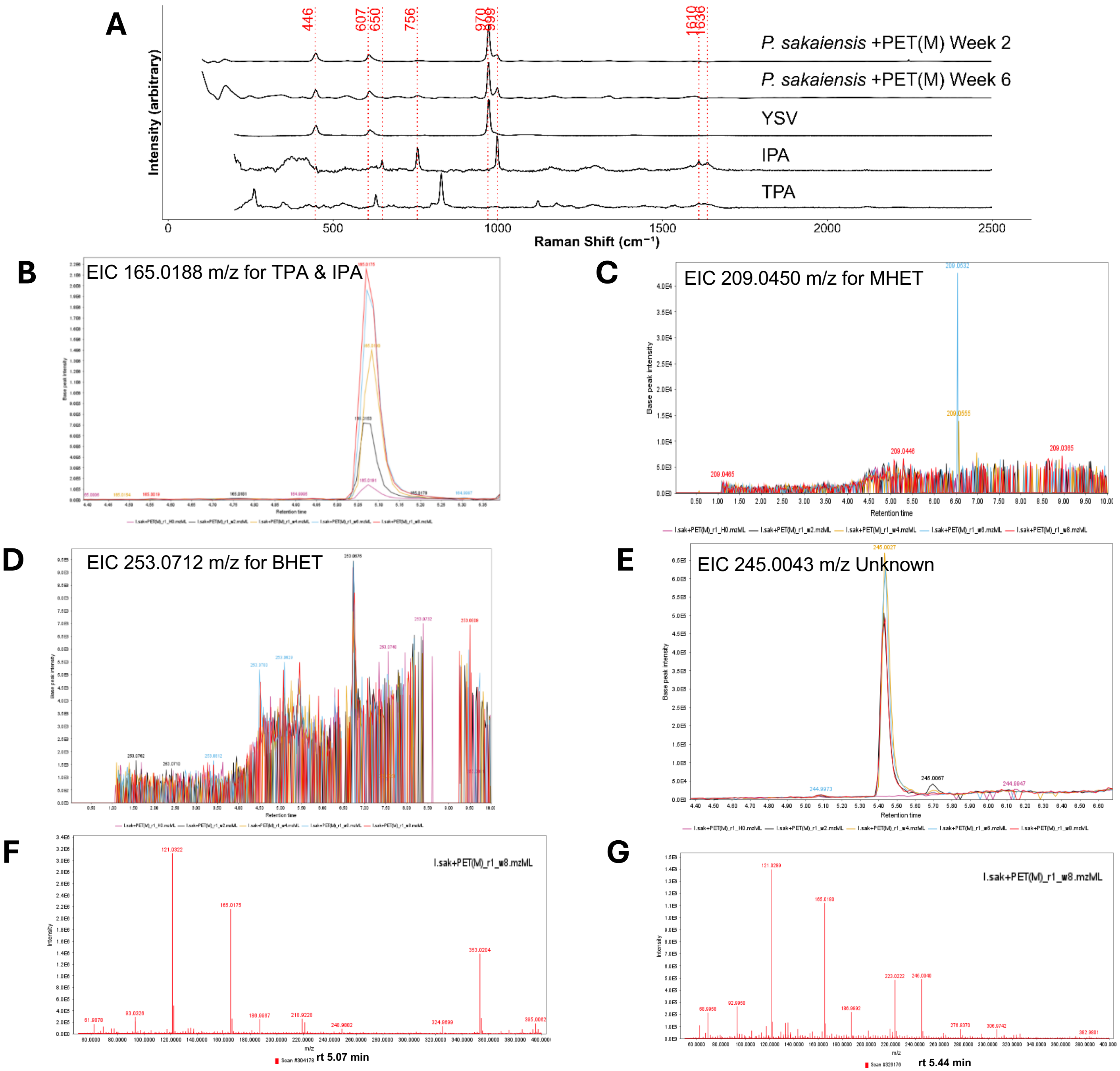

**Figure S9**: Detection of *P. sakaiensis* monomers from the metabolism of polyethylene terephthalate (PET) manufactured by Magerial Science (M) during the eight week incubation. (A) Raman spectroscopy data comparison for cell culture samples at week 2 and week 6 and isophthalic acid (IPA) and terephthalic acid (TPA) standards. (B-E) are the extracted ion chromatograms (EIC) of samples obtained from cultures from week 0 to week 8 for *m/z* 165.0188 (B), 209.0450 (C), 253.0712 (D) and 245.0043 for the unknown compound (E). (F) and (G) are the mass spectra (with in-source fragmentation) for IPA at RT of 5.07 min (G) and the unknown compound at RT of 5.44 min (G). EIC were constructed using *m/z* tolerance of 0.01 Da.

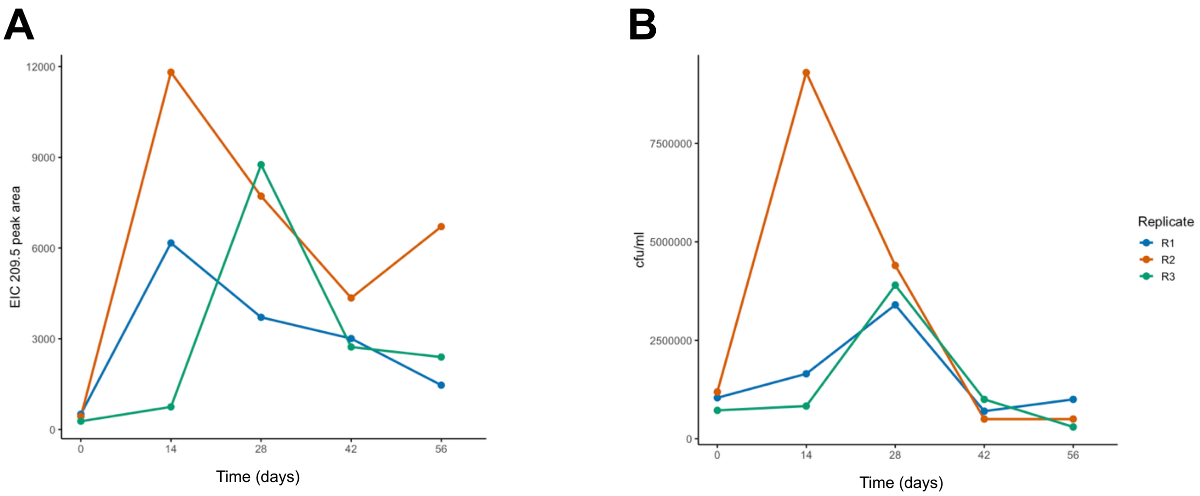

**Figure S10:** Detection of the monomer MHET during cellular growth on polyethylene terephthalate (PET) manufactured by Goodfellow during the eight-week incubation. (A) Area values for extracted ion chromatographs taken for MHET (*m/z* 209.0450) for individual cultures sampled from week 0 to week 8. (B) Cell growth for individual replicates based on viable plate counts taken from week 0 to week 8. Replicates are shown as colour coded series.

**Table S5.** Terephthalic acid (TPA) Raman spectrum band assignments from (Arenas & Marcos, 1980; Téllez S et al., 2001)

| **Raman shift ($cm^{-1}$)** | **Band assignment** |
|---|---|
| 261 | In plane bending (C=C=C) and (C=C=O) |
| 349 | Torsion (CCCC) |
| 630 | Torsion (CCCC) and (HOC=O) |
| 829 | In plane bending (C=C=C) and bending (CCC in ring) |
| 1123 | Stretching (CO) and in plane bending (COH) |
| 1179 | Stretching (CO) and in plane bending (COH) |
| 1289 | In plane bending (COH) and bending (CCH in ring) |
| 1445 | In plane bending (OH) and stretching (CO) |
| 1630 | Stretching (C=O) and in plane bending (COH) |

**Table S6**. Isophthalic acid (IPA) Raman spectrum band assignments from (Arenas & Marcos, 1980; Bardak et al., 2016)

| **Raman shift ($cm^{-1}$)** | **Band assignment** |
|---|---|
| 192 | In plane bending (CCC) and (CCO) |
| 420 | Torsion (CCCO), (CCCC), (CCOH) and (CCCH) |
| 492 | In plane bending (CCC) and (CCO) |
| 650 | Torsion (CCCC) and (CCCH) |
| 756 | Out of plane bending (OCO) |
| 827 | Out of plane bending (CH[CCCH]) and torsion (CCCO) |
| 999 | Stretching (CC in ring) and in plane bending (CCC) |
| 1167 | Stretching (CC in ring) and in plane bending (CCH) |
| 1299 | Stretching (CC in ring) and (CO) and in plane bending (COH) and (CCH) |
| 1610 | Stretching (CC in ring) and in plane bending (CCH) |
| 1636 | Stretching (C=O) |

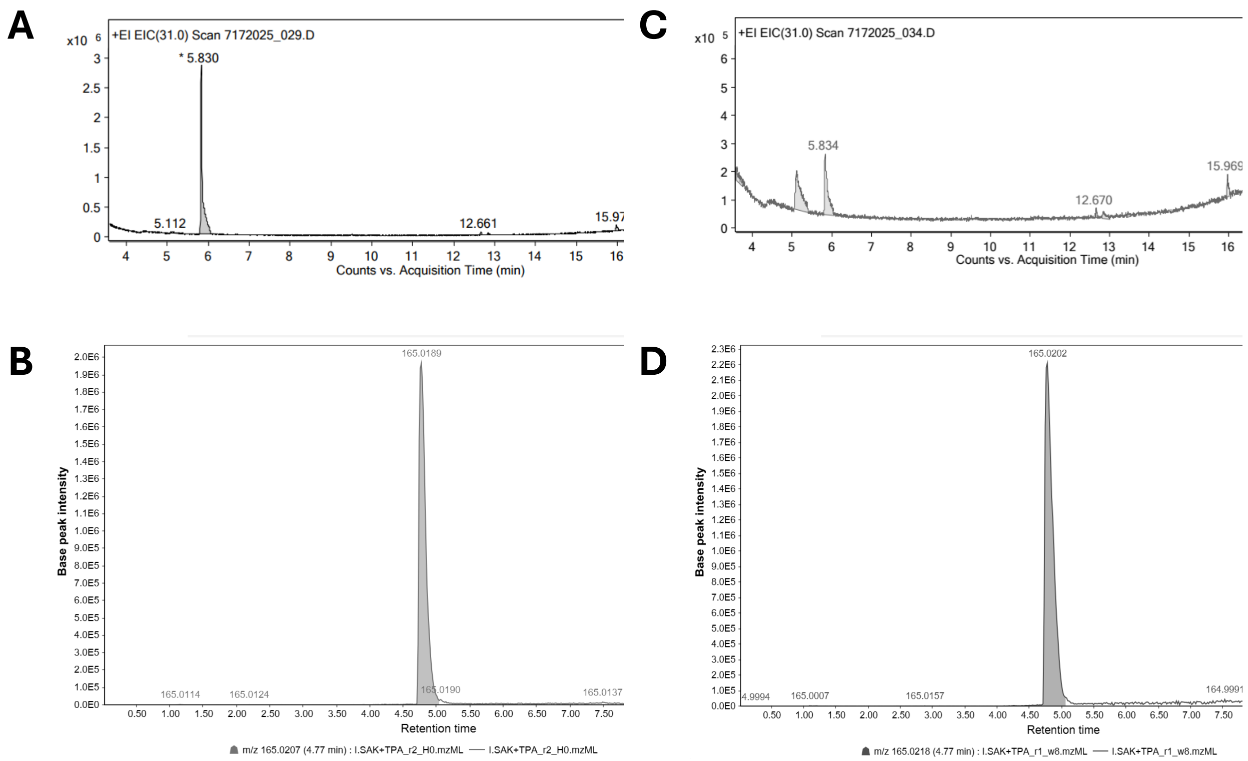

**Figure S11**: Extracted Ion Chromatograph (EIC) of *P. sakaiensis* when supplemented with expected breakdown products of polyethylene terephthalate (PET) including (A and C) ethylene glycol (EG) and (B and D) terephthalic acid (TPA) at day 0 (A and B), day 14 (C) and day 56 (D) using a 2.5-fold concentrate supernatant.

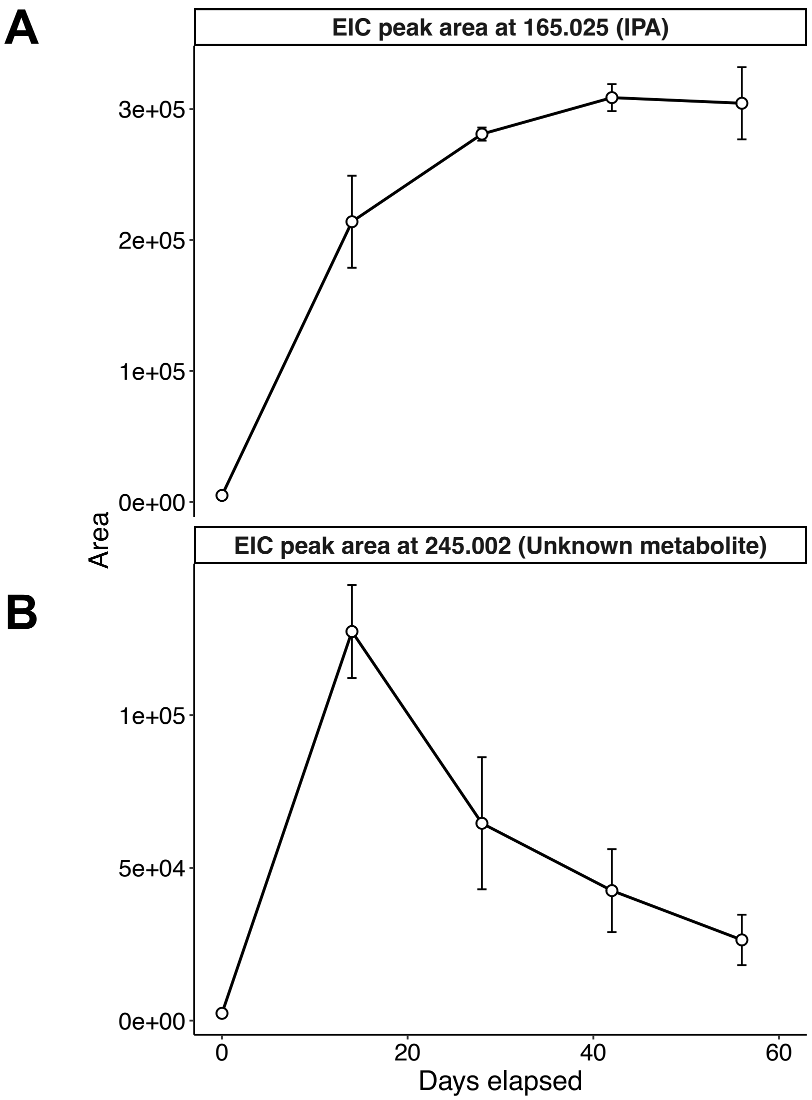

**Figure S12:** Detection of the *P. sakaiensis* monomer IPA compared to an unknown metabolite during cellular growth from the metabolism of group up polyethylene terephthalate (PET) manufactured by Goodfellow during the 8-week incubation. (A) Mean area (n = 3) values for extracted ion chromatographs (EIC) taken for IPA (*m/z* 165.025) where error bars represent standard deviation. (B) Mean area (n = 3) values for extracted ion chromatographs taken for the unknown metabolite (*m/z* 245.022) where error bars represent standard deviation.

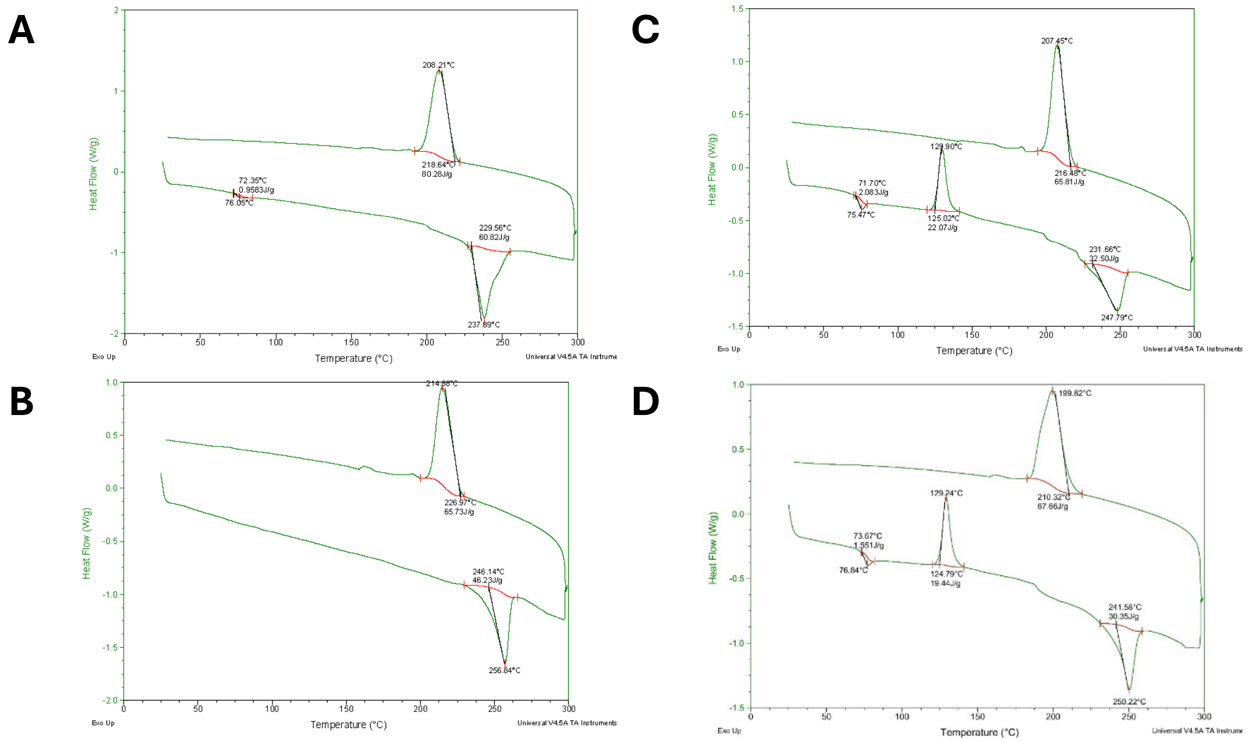

**Figure S13**: Differential scanning calorimetry (DSC) spectrums of various manufacturers and structure orientations of polyethylene terephthalate (PET) including (A) Magerial Science manufacturer ground PET powder, (B) Goodfellow manufacturer bilateral orientation PET film, (C) Goodfellow manufacturer amorphous PET film which was ground and (D) Goodfellow manufacturer amorphous PET film.

**Table S7**. Intensity signals at 1092 and 1114 $cm^{-1}$ and their ratios from Raman spectroscopy for four types of polyethylene terephthalate (PET) with varying structural orientation and crystallinity.

| | **1091.74** | **1114.74** | **Ratio** |
|---|---|---|---|
| **PET_Magerial_Grinded** | 329.75 | 187 | 1.763369 |
| **PET_Goodfellow_Amorphous_Film** | 270.5 | 464.75 | 0.582033 |
| **PET_Goodfellow_Amorphous_Grinded** | 202.5 | 327.75 | 0.617849 |
| **PET_Goodfellow_Biaxially oriented_Film** | 918.25 | 538.5 | 1.7052 |

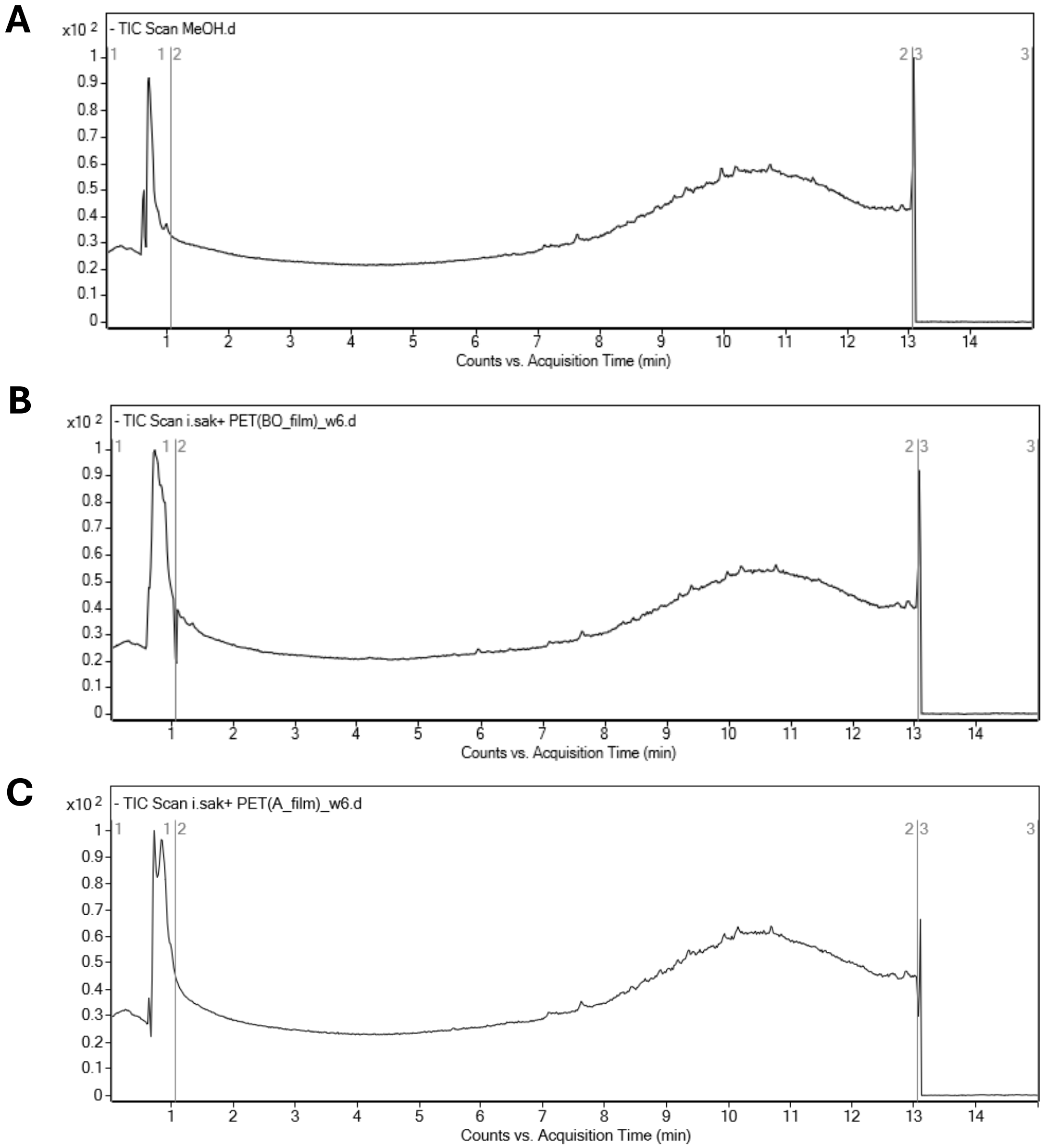

**Figure S14**: Total Ion Chromatograph (TIC) of *P. sakaiensis* supplied with various Goodfellow polyethylene terephthalate (PET) films containing different structure orientation for 6-week duration. (A) methanol blank, (B) Goodfellow biaxially orientated PET film + *P. sakaiensis* (1 cm by 1cm and 0.01 cm thick, ES30-FM-000200), and (C) Goodfellow amorphous film (1 cm by 1cm and 0.15 cm thick, ES30-SH-000115) + *P. sakaiensis*.
